# AUTOTAC-mediated targeted degradation of transthyretin aggregates ameliorates hereditary transthyretin amyloidosis

**DOI:** 10.64898/2026.02.23.707350

**Authors:** Hee Yeon Kim, Daniel Youngjae Park, Eun Hye Cho, Yeon Sung Son, Sung Hyun Kim, Ki Woon Sung, Helena Sofia Martins, Maria João Saraiva, Maria Rosário Almeida, Chang Hoon Ji, Yong Tae Kwon

## Abstract

Hereditary transthyretin amyloidosis (hATTR) is characterized by extracellular deposition of amyloidogenic transthyretin (TTR) aggregates, yet the mechanisms governing their clearance remains poorly understood. Here, we identify a key role for the N-degron pathway in lysosomal degradation of the pathogenic TTR^V30M^ variant. Misfolded intracellular TTR^V30M^ was rapidly secreted and subsequently re-entered within 24 hours during cell-to-cell trafficking. The molecular chaperone R-BiP—N-terminally (Nt) arginylated HSPA5/BiP/GRP78— associated with intracellular TTR^V30M^, and its Nt-arginine functioned as an agonist for the N-recognin sequestosome 1 (SQSTM1/p62). This interaction facilitated p62-dependent autophagosomal sequestration and lysosomal degradation of TTR^V30M^. To pharmacologically exploit this mechanism, we applied the AUTOTAC (AUTOphagy-TArgeting Chimera) platform, which enables the targeting of substrates to p62 for autophagic clearance. We developed Autotac 201 (ATC201), an 876-Da chimera designed to bind both the T4 pocket of aggregated TTR and p62, thereby promoting selective autophagic degradation. In cultured cells, ATC201 potently reduced intracellular TTR^V30M^ aggregates in a manner depending on p62-mediated autophagy, exhibiting a DC₅₀ of low nM. In hATTR model mice, ATC201 markedly lowered tissue TTR aggregate burden and restored autophagy pathway flux impaired by aggregate accumulation. Treatment improved nerve conduction parameters and reduced peripheral neuropathy scores, indicating functional rescue. ATC201 also led to preservation of muscle strength and attenuation of systemic amyloid deposition. Our findings reveal that the N-degron pathway orchestrates autophagic removal of TTR aggregates and demonstrate the therapeutic potential of AUTOTAC-based degraders for hATTR and other proteinopathies characterized by pathogenic protein aggregation.

## Introduction

Proteinopathies comprise a diverse group of disorders characterized by the deposition of misfolded proteins that assemble into highly ordered fibrillary structures [1]. These pathogenic agents can accumulate intracellularly, as exemplified by tau in Alzheimer disease, α-synuclein in Parkinson disease, and TDP-43 in amyotrophic lateral sclerosis [2, 3]. In contrast, aggregates such as amyloid-β in Alzheimer disease and transthyretin (TTR) in hereditary transthyretin amyloidosis (hATTR) predominantly accumulate in the extracellular milieu, where they exert their pathogenic effects [4, 5]. Some aggregates, including α-synuclein, also exhibit prion-like properties, acting as seeding particles that propagate between cells and drive the progressive spread of pathology [6]. Both intracellular and extracellular aggregates promote the formation of higher-order fibrillar assembles, amplifying proteotoxic stress and contributing to disease progression [1].

Caused by pathogenic variants in the TTR gene, hATTR is a fatal systemic proteinopathy [7]. TTR is a 127-amino-acid extracellular protein that normally forms a stable homotetramer functioning as a carrier for retinol binding protein and thyroxine (T4). In hATTR, pathogenic mutations destabilize the tetramer, promoting its dissociation into misfolded monomers that assemble into amyloid fibrils [8]. These fibrils progressively deposit across multiple organs—including peripheral nerves, the heart, kidneys, and ocular tissues—reflecting systemic dissemination of the aggregated species [9–11]. Among more than 140 pathogenic mutations, the most common are V30M and V122I, which underlie familial amyloid polyneuropathy (FAP) and familial amyloid cardiomyopathy, respectively [12, 13]. Clinically, hATTR manifests progressive polyneuropathy, autonomic dysfunction, and restrictive cardiomyopathy that together may culminate in severe disability and premature death within a decade of symptom onset [14]. Beyond hereditary disease, wild-type TTR amyloidosis (ATTR), formerly called senile systemic amyloidosis, arises without genetic mutations. Age-related destabilization of wild-type TTR promotes misfolding and fibril deposition predominantly in the myocardium, causing restrictive cardiomyopathy in elderly individuals [15, 16]. Autopsy studies revealed myocardial ATTR deposits in ∼25% of individuals aged ≥85 years, and prospective screening detected cardiac ATTR in 10–16% of patients with severe aortic stenosis [17, 18]. Both hereditary and wild-type forms remain substantially undiagnosed, suggesting that the true incidence exceeds recognized cases [19].

Current therapies include tetramer stabilizers such as tafamidis or RNA interference–based agents including patisiran and inotersen [20, 21]. Although these treatments slow disease progression, they do not eliminate pre-existing aggregates [22, 23]. This therapeutic gap underscores the critical need for disease-modifying therapeutic capable of removing pathogenic TTR species and their accumulated deposits.

Cells employ protein quality control systems to eliminate misfolded proteins, prominently the ubiquitin (Ub)-proteasome system (UPS) and autophagy-lysosome system (hereafter autophagy) [24]. The UPS serves as the first-line mechanism for degrading soluble misfolded proteins [25]. Misfolded proteins that escape UPS-mediated degradation or are prone to forming aggregates are directed to macroautophagy/autophagy [26], where they are sequestered into autophagosomes that subsequently fuse with lysosomes for degradation [27]. Autophagic cargo recognition is mediated by selective receptors such as sequestosome 1 (SQSTM1/p62) bind ubiquitinated or tagged substrates and facilitate their co-degradation by lysosomal hydrolases [28, 29]. Proteinopathies arise when misfolded proteins accumulate due to resistance or overload of both the UPS and autophagy [30, 31]. Although the cellular mechanisms governing the clearance of intracellular protein aggregates have been extensively studied, relatively little is known about how extracellular protein aggregates are recognized and removed by intracellular quality control systems.

The N-degron pathway regulates the degradation of a wide spectrum of proteins and biomaterials, including subcellular organelles and invading pathogens [32, 33]. In this system, single Nt amino acids and their structural derivatives—collectively termed N-degrons—are recognized by N-recognins that couple substrates to either the UPS or autophagy [34]. In the UPS, known N-degrons include Arg, Lys, and His (type 1, positively charged) as well as Trp, Phe, Tyr, Leu, and Ile (type 2, bulky hydrophobic) [35, 36]. These degrons are bound by the UBR box of N-recognins such as UBR1, UBR2, UBR4, and UBR5, leading to substrate ubiquitination and proteasomal degradation [33]. Amongst these, Nt-Arg can be generated post-translationally by arginyl-tRNA–protein transferase 1 (ATE1), which conjugates L-Arg to defined Nt-residues [26, 37, 38]. We have previously demonstrated that N-degrons also mediate p62-dependent autophagy [29, 39]. Upon accumulation of protein aggregates or other UPS-resistant cargoes, a subset of endoplasmic reticulum (ER)-resident chaperones, including GRP78/BiP, undergo retrotranslocation and Nt-arginylation [29, 40]. The resulting Arg/N-degrons bind by the ZZ-type zinc finger (ZZ) domain of the N-recognin p62, also known as an autophagic receptor, allosterically activating p62 to self-polymerize and form high-molecular-weight condensates via liquid-liquid phase separation [39, 41].

Concurrently, exposure LC3-interacting region (LIR) enables engagement with LC3-II on phagophores, promoting sequestration of p62-cargo complexes into autophagosomes and subsequent lysosomal degradation [42, 43]. Building on this allosteric mechanism, we developed synthetic mimetics of Arg/N-degrons—autophagy-targeting ligands (ATLs) —that pharmacologically enhance autophagic degradation of diverse biomaterials, including protein aggregates [26, 44], subcellular organelles [45–48], and invading pathogens [49].

Targeted protein degradation (TPD) has emerged as a transformative paradigm in drug development due to its ability to eliminate proteins previously considered undruggable [50]. Amongst TPD technologies, proteolysis-targeting chimeras (PROTACs) recruit E3 ubiquitin ligases to induce ubiquitination and proteasomal degradation of target proteins [51]. Although extensively investigated—particularly for oncogenic drivers— PROTACs face several limitations, including off-target effects and the emergence of drug resistance through mutations that impair E3 ligase engagement or substrate ubiquitination [52]. Moreover, because the proteasome cannot unfold protein aggregates through its restricted chamber, it remains unclear how proteasome-based TPD approaches can effectively address aggregated or fibrillar species [53–55]. To overcome these challenges, we developed AUTOTAC (AUTOphagy-TArgeting Chimera), an autophagy-based TPD platform designed to degrade a broader range of pathogenic substrates, including aggregates. AUTOTACs are bifunctional molecules consisting of a target-binding ligand (TBL) linked to an ATL, a chemical N-degron that activates the p62 ZZ domain through the autophagic N-degron pathway [56]. This strategy has been validated proof-of-concept studies demonstrating the targeted degradation of α-synuclein aggregates in Parkinson disease [44] and the androgen receptor splice variant AR-V7 in castration-resistant prostate cancer [57].

This study was designed to develop a therapeutic strategy for hATTR by targeting pathogenic TTR aggregates. We found that misfolded TTR aggregates were rapidly secreted and subsequently re-entered cells within 24 hours, indicating a dynamic cycle of extracellular dissemination and re-internalization. Intracellular pathogenic TTR species associated with R-BiP, which allosterically activated p62 as an N-recognin and mediated lysosomal co-degradation of the aggregates. ATC201, a bifunctional degrader engineered to direct pathogenic TTR aggregates to p62, robustly induced their clearance in cultured cells, exhibiting DC_50_ and D_max_ values of low nM. In hATTR mouse models, ATC201 significantly reduced TTR aggregate burden across multiple organs, including the colon, small intestine, liver, brain, and muscle. The extent of aggregate degradation correlated with marked improvements in systemic and neuromuscular function. These results uncover a previously uncharacterized role of the N-degron pathway in the quality control of pathogenic TTR aggregates and support AUTOTAC as a promising therapeutic strategy for systemic amyloidosis, with broader implications for treating proteinopathies driven by aggregate accumulation.

## Results

### Cell-to-cell transmission of amyloidogenic TTR and its impact on ER stress

To date, little is known about the mechanism by which extracellular pathogenic TTR species contribute to hATTR pathogenesis in distant tissues. To address this, we monitored the metabolic fates of two amyloidogenic variants, TTR^V30M-MYC^ and TTR^V122I-MYC^, compared with wild-type (WT) TTR, following transient expression in HeLa and HEK293T cells, both of which lack endogenous TTR. Thioflavin-T (ThT) fluorescence measurements revealed that intracellular TTR^V30M^ readily formed amyloid fibrils (Fig. 1A, 1B and S1A), consistent with the high aggregation-prone propensity of its monomeric form (Fig. 1C). Misfolded TTR^V30M^ monomers readily assembled into oligomers (Fig. 1D), of which 40-60% were subsequently secreted into the extracellular medium (Fig. 1E). When recipient cells were incubated with conditioned media (CM) containing secreted TTR variants (Fig. 1F), up to 30% of total TTR species present in CM were recovered from cellular extracts (Fig. 1G and S1B), indicating efficient uptake. Internalized TTR species formed punctate cytosolic structures (Fig. 1H, 1I, S1C, S1D and S1E) as early as 3 hours post-exposure, reaching maximal levels at 18 hours (Fig. 1H and 1I). These results suggest that misfolded TTR variants rapidly form oligomers and aggregates both inside and outside the cell and are subsequently transmitted between cells. Importantly, both extracellular and intracellular TTR^V30M^ species triggered ER stress as evidenced by increases of the protein level of CHOP (Fig. 1G and 1J) and the mRNA levels of ATF4, spliced XBP1, and CHOP (Fig. 1K). Likewise, the ER stress inducer thapsigargin markedly accelerated the aggregation of TTR^V30M^ (Fig. 1L). These results suggest that amyloidogenic TTR mutants and ER stress amplify one another in a pathological feedback loop that may drive the progression of hATTR.

**Figure 1.**
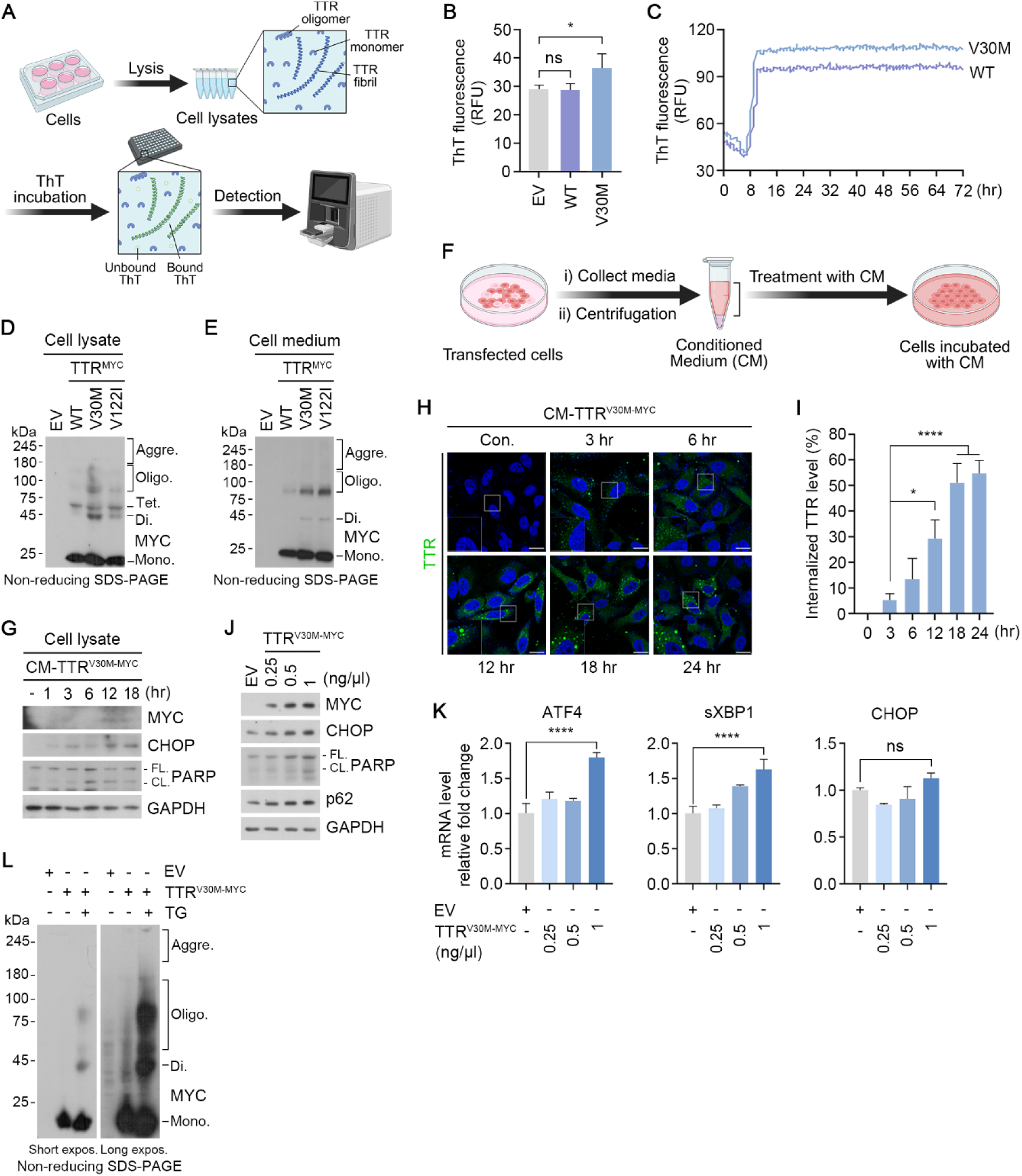
Amyloidogenic TTR exhibits propagation-prone behavior and disrupts proteostasis. (**A**) Schematic illustration of thioflavin-T (ThT) assay with cell lysates. (**B**) Quantification of ThT fluorescence of lysates from HeLa cells transfected with WT TTR^MYC^ or TTR^V30M-MYC^ (n=3). (**C**) ThT assay with purified Myc-tagged WT TTR and TTR^V30M^ proteins from HEK293T cells. (**D**) *In vivo* oligomerization assay of lysates from in HeLa cells transfected with TTR variants. (**E**) *In vivo* oligomerization assay of TCA-precipitated media from HeLa cells transfected with TTR variants. (**F**) Schematic illustration of the experimental setup using conditioned media (CM) containing secreted TTR variants. (**G**) Immunoblotting analysis of HeLa cells treated with CM containing secreted TTR^V30M-MYC^ at the indicated time points. (**H**) Immunostaining analysis of HeLa cells treated with CM containing secreted TTR^V30M-MYC^ at the indicated time points. (**I**) Quantification of (H) (n = 50 cells). (**J**) Immunoblotting analysis of HeLa cells transfected with TTR^V30M-MYC^ at the indicated concentrations. (**K**) The mRNA expression relative to *GAPDH* was measured by RT-qPCR. RNAs were isolated from HeLa cells transfected with TTR^V30M-MYC^ at the indicated concentrations (n=3). (**L**) *In vivo* oligomerization assay of HeLa cells transfected with TTR^V30M-MYC,^ subsequently treated with thapsigargin (200 nM, 18 h). Scale bar represents 10μm. Data are presented as means ± SD. Each n represents an independent biological replicate. Unpaired two-tailed Student’s *t*-test (B); one-way ANOVA (I, K); p* < 0.05, **p < 0.01, ***p < 0.001, ****p < 0.0001 and ns, not significant.

## The ER-residing molecular chaperone BiP senses and counteracts amyloidogenic TTR as part of protein quality control

To elucidate how pathogenic TTR species are recognized and handled by protein quality control machinery, we screened a panel of molecular chaperones known to associate with TTR [58, 59]. We noted that the chaperone BiP, retrotranslocated from the ER, was upregulated at both the protein (Fig. 2A) and mRNA levels (Fig. 2B) following expression of TTR^V30M^. Co-immunoprecipitation (co-IP) assays demonstrated that BiP associated with TTR^V30M^, and this interaction was further strengthened by ER stress (Fig. 2E and S2A).

**Figure 2.**
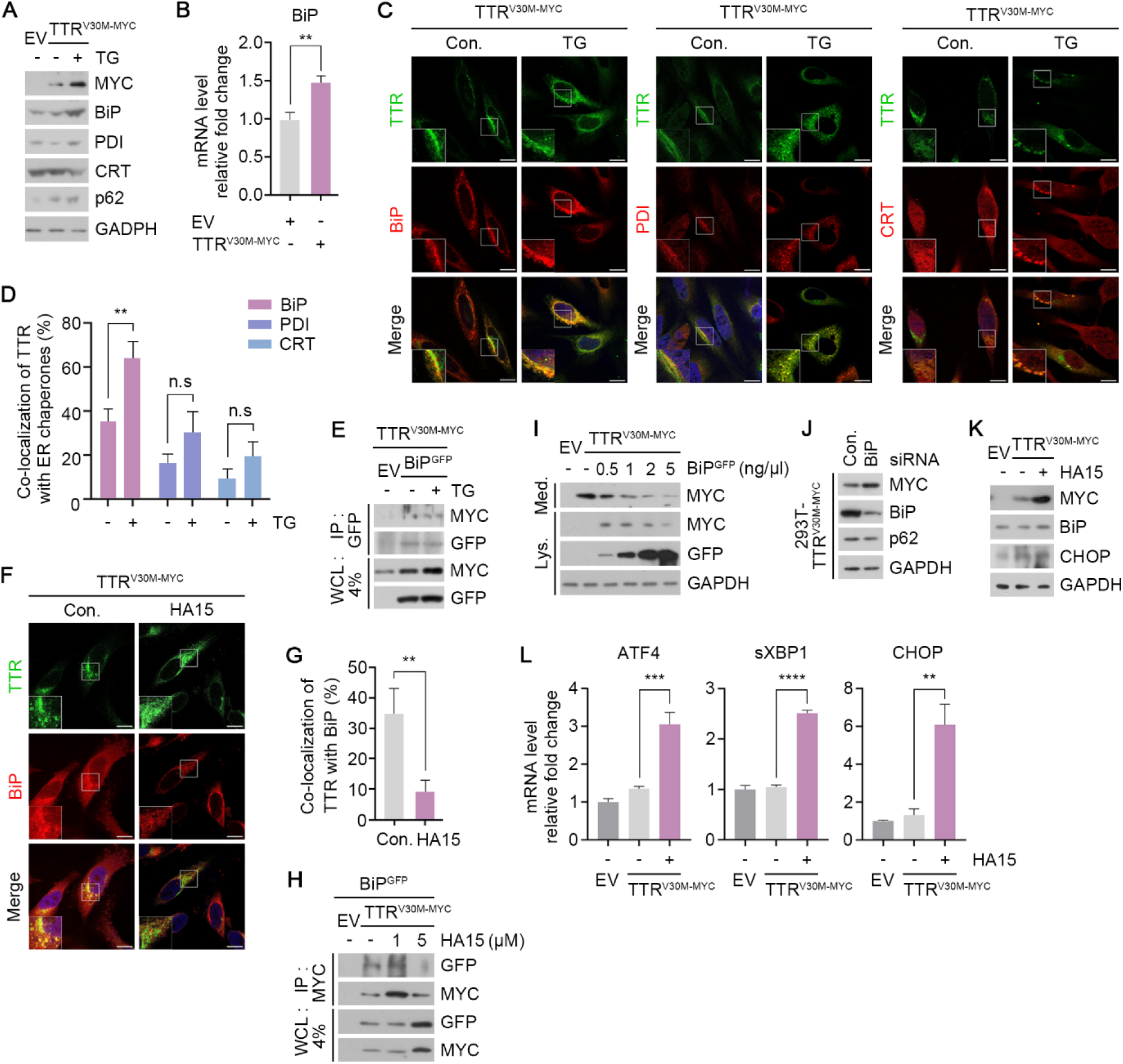
The ER-resident molecular chaperone BiP senses and regulates amyloidogenic TTR^V30M^. (**A**) Immunoblotting analysis of HeLa cells transfected with TTR^V30M^ treated with thapsigargin (200 nM, 18 h). (**B**) The mRNA expression relative to GAPDH was measured by RT-qPCR. RNAs were isolated from HeLa cells transfected with TTR^V30M-MYC^ (n=3). (**C**) Immunostaining analysis of HeLa cells transfected with TTR^V30M-MYC^, subsequently treated with thapsigargin (200 nM, 18 h). (**D**) Quantification of (C) (n = 50 cells). (**E**) CoIP assay of HEK293T cells co-transfected with TTR^V30M-MYC^ and BiP ^GFP^, subsequently treated with thapsigarin (200 nM, 18 h). (**F**) Immunostaining analysis of HeLa cells transfected with TTR^V30M-MYC^, subsequently treated with HA15 (5 μM, 24 h). (**G**) Quantification of (F) (n = 50 cells). (**H**) CoIP assay of HEK293T cells co-transfected with TTR^V30M-MYC^ and BiP ^GFP^, subsequently treated with HA15 at the indicated concentrations (24 h). (**I**) Immunoblotting analysis of lysates and TCA-precipitated media from HeLa cells co-transfected with TTR^V30M-MYC^ and BiP^GFP^ at the indicated concentrations (24 h). (**J**) Immunoblotting analysis of TTR^V30M-MYC^ expressing 293T stable cell line transfected with siRNAs. (**K**) Immunoblotting analysis of HeLa cells transfected with TTR^V30M-MYC^, subsequently treated with HA15 (5 μM, 24 h). (**L**) The mRNA expression relative to GAPDH was measured by RT-qPCR. RNAs were isolated from HEK293T cells transfected with TTR^V30M-MYC^, subsequently treated with HA15 (5 μM, 24 h) (n=3). Scale bar represents 10μm. Data are presented as means ± SD. Each n represents an independent biological replicate. Unpaired two-tailed Student’s t test; p* < 0.05, **p < 0.01, ***p < 0.001, ****p < 0.0001 and ns, not significant.

Consistent with a functional association, immunostaining revealed cytosolic puncta doubly positive for BiP and TTR^V30M^ (Fig. 2C and 2D). Their interaction (Fig. 2H) and puncta formation (Fig. 2F and 2G) were impaired by HA15 that inhibits the ATPase activity of BiP. The level of BiP exhibited an inverse correlation with t intracellular and extracellular TTR^V30M^ abundance (Fig. 2I and 2J). Inhibition of BiP ATPase activity with HA15 lead to the accumulation of TTR^V30M^ (Fig. 2K), which in turn provoked ER stress (Fig. 2L). These results indicate that cytosolic BiP—retrotranslocated from the ER—functions as a sensor for amyloidogenic TTR and counteracts its intracellular and extracellular accumulation, which requires the ATPase activity of BiP.

### Nt-arginylation of BiP promotes lysosomal degradation of amyloidogenic TTR

Misfolded TTR species are generally thought to be eliminated primarily through the UPS [59]. However, in HeLa cells expressing TTR^V30M^, we found that degradative flux was unaffected by the proteasomal inhibitor MG132 but was strongly blocked by the autophagy inhibitor hydroxychloroquine (HCQ) (Fig. 3A). Consistently, ATG5 knockdown led to TTR accumulation across multiple cell lines (Fig. S3A and S3B), indicating that autophagy, rather than the UPS, mediates degradation of amyloidogenic TTR under these conditions. We next tested whether this autophagic degradation requires the Arg/N-degron on BiP. which can be post-translationally conjugated by ATE1-mediated Nt-arginylation. Using a newly generated antibody specific to R-BiP, we observed an inverse correlation between R-BiP and TTR^V30M^ levels (Fig. 3B). To further probe the role of Nt-arginylation, we employed an Ub fusion technique, in which Ub-X-BiP^GFP^ was co-translationally cleaved by deubiquitinases to generate X-BiP^GFP^ bearing defined Nt-residues (X= R, E or V) (Fig. 3C). Immunoblotting revealed that R-BiP^GFP^, but not the E- or V-bearing variants, induced robust degradation of TTR^V30M^ (Fig. 3D). This activity was abolished HCQ treatment (Fig. 3E), demonstrating that R-BiP promotes lysosomal degradation of TTR^V30M^ species.

**Figure 3.**
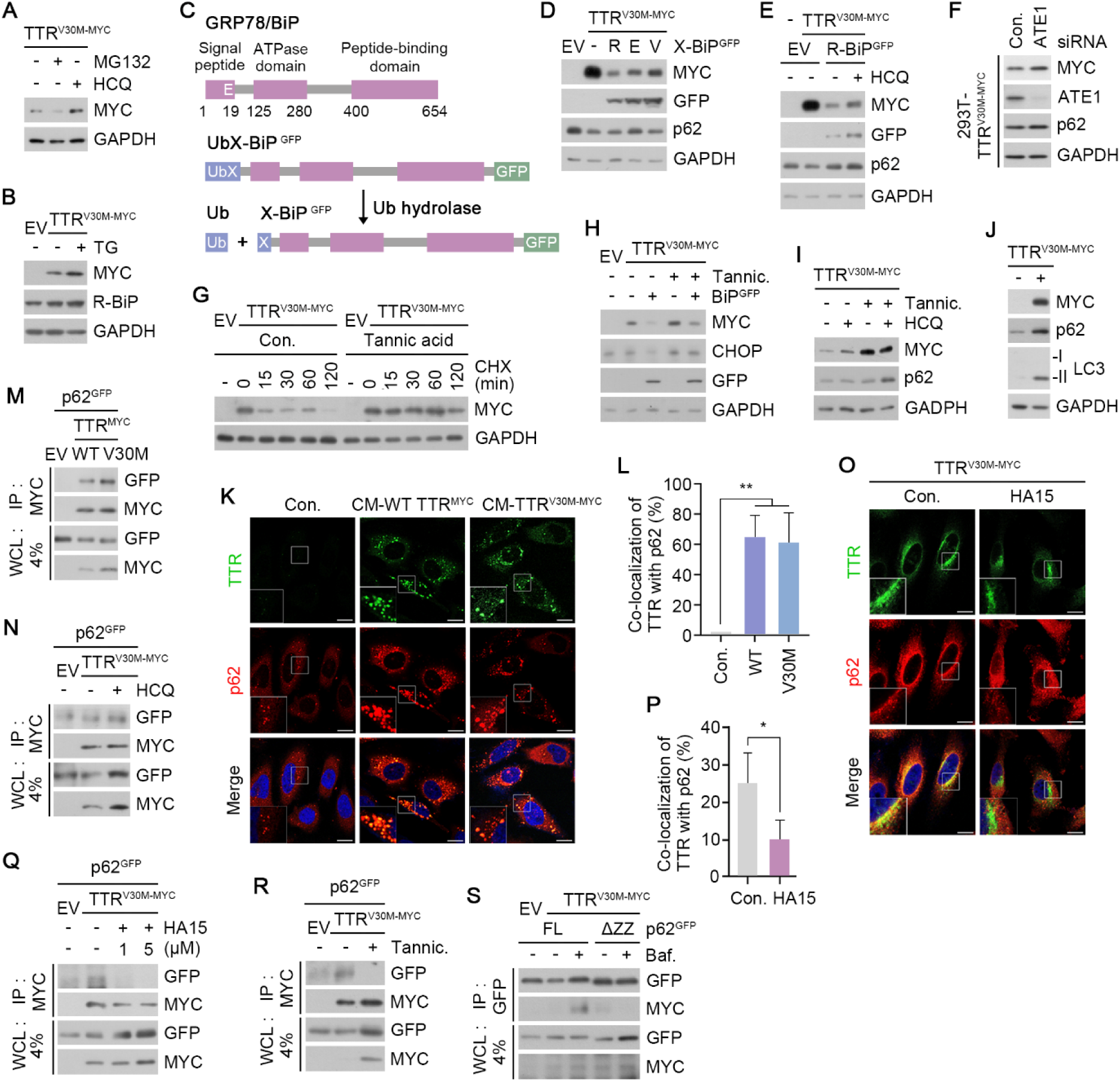
Nt-arginylated BiP forms a complex with p62 and TTR for autophagic degradation. (**A**) Immunoblotting analysis of HeLa cells transfected with TTR^V30M-MYC^, subsequently treated with MG132 (2 μM, 18h) or HCQ (10 μM, 24h). (**B**) Immunoblotting analysis of HeLa cells transfected with TTR^V30M-MYC^, subsequently treated with thapsigargin (200 nM, 18 h). (**C**) Schematic illustration of full length BiP in comparison with Ub-X-BiP ^GFP^, in which Ub is C-terminally conjugated with N-terminal X residues (X= R, E or V) of BiP ^GFP^. (**D**) Immunoblotting analysis of HeLa cells co-transfected with TTR^V30M-MYC^ and Ub-X-BiP ^GFP^ (X= R, E or V). (**E**) Immunoblotting analysis of HeLa cells co-transfected with TTR^V30M-MYC^ and R-BiP ^GFP^, subsequently treated with HCQ (10 μM, 24h). (**F**) Immunoblotting analysis of TTR^V30M-MYC^ expressing 293T stable cell line transfected with siRNAs. (**G**) Cycloheximide (CHX) chase assay of HeLa cells transfected with TTR^V30M-MYC^ under tannic acid (25 μM, 24 h) treatment. (**H**) Immunoblotting analysis of HeLa cells co-transfected with TTR^V30M-MYC^ and BiP ^GFP^, subsequently treated with tannic acid (25 μM, 24 h). (**I**) Immunoblotting analysis of HeLa cells transfected with TTR^V30M-MYC^, subsequently treated with tannic acid (25 μM, 24 h) in the presence or absence of HCQ (10 μM, 24 h). (**J**) Immunoblotting analysis of HeLa cells transfected with TTR^V30M-MYC^. (**K**) Immunostaining analysis of HeLa cells treated with CM containing secreted WT TTR^MYC^ or TTR^V30M-MYC^ for 24 h. (**L**) Quantification of (K) (n = 50 cells). (**M**) CoIP assay in HEK293T cells co-transfected with p62^GFP^ and TTR^MYC^. (**N**) CoIP assay in HEK293T cells co-transfected with p62^GFP^ and TTR^V30M-MYC^, subsequently treated with HCQ (10 μM, 24h). (**O**) Immunostaining analysis of HeLa cells transfected with TTR^V30M-MYC^, subsequently treated with HA15 (5 μM, 24 h). (**P**) Quantification of (O) (n = 50 cells). (**Q**) CoIP assay in HEK293T cells co-transfected with p62^GFP^ and TTR^V30M-MYC^, subsequently treated with HA15 at the indicated concentrations (24 h). (**R**) CoIP assay in HEK293T cells co-transfected with p62^GFP^ and TTR^V30M-MYC^, subsequently treated with tannic acid (25 μM, 24 h). (**S**) CoIP assay in HEK293T cells co-transfected with p62^GFP^ (Full length or ZZ domain deleted) and TTR^V30M-MYC^, subsequently treated with Bafilomycin A1 (50 nM, 3h). Scale bar represents 10μm. Data are presented as means ± SD. Each n represents an independent biological replicate. Unpaired two-tailed Student’s *t*-test (P); one-way ANOVA (L); p* < 0.05, **p < 0.01, ***p < 0.001, ****p < 0.0001 and ns, not significant.

We next examined the requirement of ATE1 in this process. Genetic depletion of ATE1 markedly stabilized TTR^V30M^ (Fig. 3 F), and pharmacological inhibition of ATE2 with tannic acid extended the half-life of TTR^V30M^ from ∼40 min to over 120 min (Fig. 3G). Non-reducing SDS-PAGE, performed to monitor oligomeric states, revealed dramatic accumulation of both monomeric and high-molecular weight TTR^V30M^ species following ATE1 inhibition (Fig. S3C). Moreover, ATE1 inhibition not only impaired TTR degradation but also suppressed the ability of BiP to mitigate ER stress (Fig. 3H). This accumulation was accompanied by reduced flux of TTR^V30M^ toward the lysosome (Fig. 3I), further supporting a role for R-BiP in mediating autophagic degradation of pathological TTR.

## R-BiP functions as a molecular bridge that recruits p62 to pathological TTR for autophagic degradation

Given our previous findings that the Nt-Arg residue of R-BiP binds and conformationally activates the autophagic receptor p62 via its ZZ domain [29, 45, 56], we next investigated whether p62 mediates TTR^V30M^ degradation. Immunoblotting showed that ectopic expression of TTR^V30M^ or incubation with CM containing extracellular TTR^V30M^ increased levels of p62 and lipidated LC3 (Fig. 3J and S3D). In immunofluorescent analyses, extracellular TTR^V30M^ in CM was internalized and formed p62-TTR colocalizing puncta throughout the cytosol (Fig. 3K and 3L), with similar results observed upon incubation with purified TTR ^V30M-MYC^ (Fig. S3E). Consistent with these observations, co-IP assays demonstrated a robust interaction between p62 and TTR^V30M^ (Fig. 3M), and degradation of p62-TTR complexes was inhibited by HCQ, indicating co-degradation via lysosome (Fig. 3N).

We then examined whether R-BiP acts as a molecular bridge between pathological TTRs and p62. Immunostaining (Fig. 3O and 3P) and co-IP assays (Fig. 3Q) revealed that the interaction between p62 and TTR^V30M^ was disrupted by HA15 treatment, suggesting that the ATPase activity of BiP is crucial for this interaction. In addition, the p62-TTR^V30M^ interaction was disrupted by the enzymatic inhibition of ATE1 using tannic acid (Fig. 3R), indicating that Nt-arginylation of BiP is required to bridge p62 to pathological TTR. Furthermore, deletion of the ZZ domain in p62 (Fig. S3F) abrogated its interaction with TTR^V30M^ and impaired autophagic degradation of the complex (Fig. 3S). These results suggest that misfolded TTRs are targeted for lysosomal degradation by the N-degron pathway, in which R-BiP acts as a link between p62 and TTRs.

## Chemical mimetics of Nt-Arg activates autophagy without effective degradation of pathological TTR variants

Currently, no therapeutic strategy efficiently removes pathogenic mutant TTR aggregates [20, 21]. We therefore examined whether mutant TTR aggregates could be degraded by autophagy-targeting ligands (ATLs), chemical N-degron memetics that we previously developed [39, 56]. To enhance the autophagy-activating efficacy, prototype compounds were further optimized through structure–activity relationship studies, resulting in identification of two compounds, ATL8 and ATL30 (Fig. 4A). We first assessed the activities of ATLs to induce p62 self-oligomerization as well as LC3 synthesis and lipidation, thereby enhancing autophagic degradation. *In vitro* oligomerization assay confirmed that ATLs efficiently induced self-polymerization of p62, a crucial step for cargo condensation (Fig. 4B). In immunocytochemistry, ATL treatment accelerated p62 puncta formation (Fig. 4C and 4D), most of which colocalized with LC3^+^ autophagic membranes (Fig. 4E and 4F), indicative of increased p62-dependent autophagy. Consistently, ATLs enhanced p62 and LC3 levels in immunoblotting analyses (Fig. 4G). These effects were also observed in cells transiently or stably expressing TTR^V30M-MYC^ (Fig. S4A, S4B and S4C).

**Figure 4.**
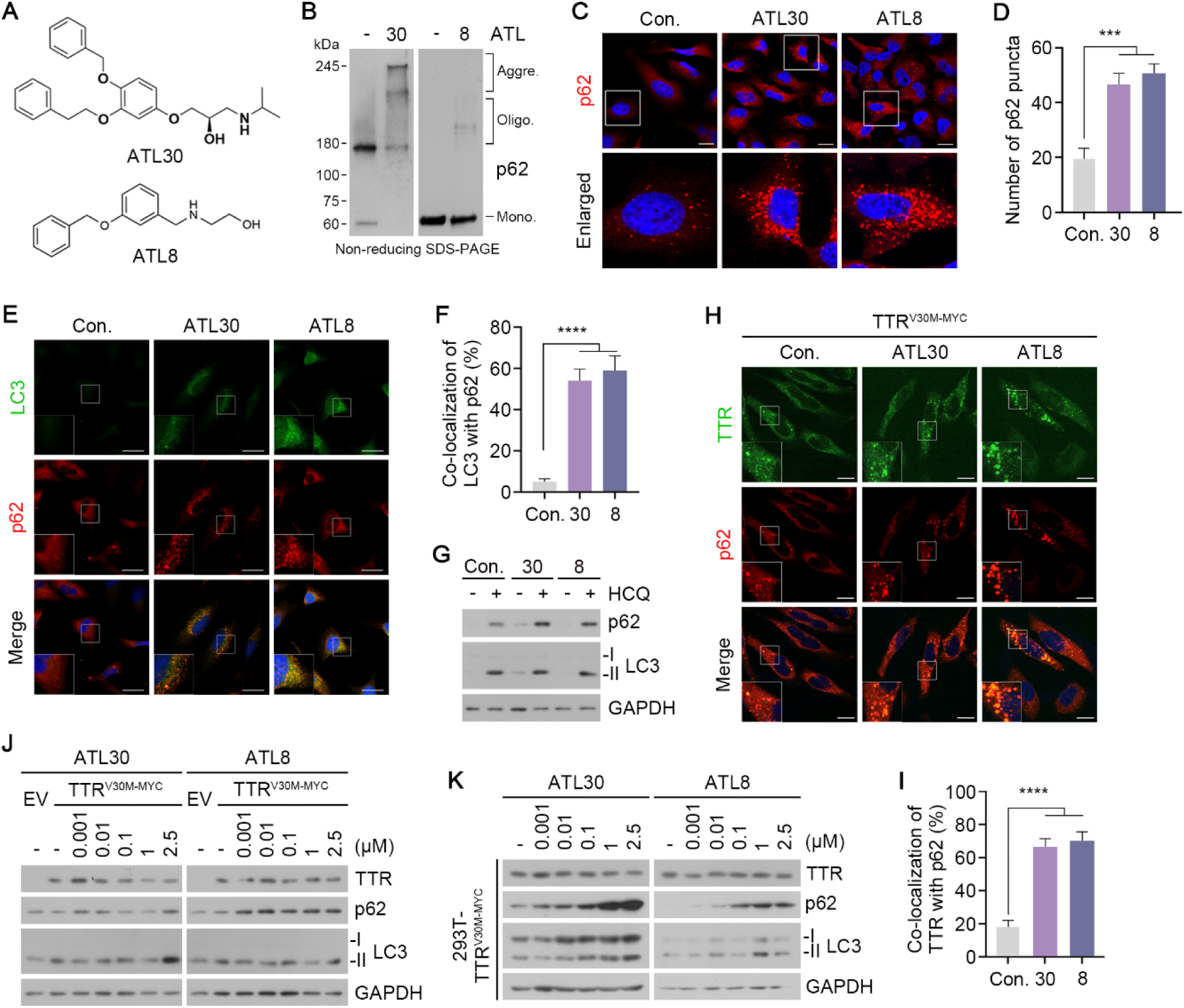
Activation of p62-dependent autophagy counteracts pathological TTR accumulation. (**A**) Chemical structures of autophagy-targeting ligands, ATL30 and ATL8. (**B**) *In vitro* p62 oligomerization assay in HEK293T cells incubated with ATL30 or ATL8 (1 μM, 24 h). (**C**) Immunostaining analysis of HeLa cells treated with ATL30 or ATL8 (1 μM, 24 h). (**D**) Quantification of (C) (n = 50 cells). (**E**) Identical to (C). (**F**) Quantification of (E) (n = 50 cells). (**G**) Immunoblotting analysis of HeLa cells treated with ATL30 or ATL8 (2.5 μM, 24 h) in the presence or absence of HCQ (10 µM, 24 h). (**H**) Immunostaining analysis of HeLa cells transfected with TTR^V30M-MYC^, subsequently treated with ATL30 or ATL8 (1 μM, 24 h). (**I**) Quantification of (H) (n = 50 cells). (**J**) Immunoblotting analysis of HeLa cells transfected with TTR^V30M-MYC^, subsequently treated with ATL30 or ATL8 at the indicated concentrations (24 h). (**K**) Immunoblotting analysis of TTR^V30M-MYC^ expressing 293T stable cell line treated with ATL30 or ATL8 at the indicated concentrations (24 h). Scale bar represents 10μm. Data are presented as means ± SD. Each n represents an independent biological replicate. One-way ANOVA; p* < 0.05, **p < 0.01, ***p < 0.001, ****p < 0.0001 and ns, not significant.

Next, we determined whether ATLs could promote degradation of TTR^V30M-MYC^. Immunocytochemistry revealed that ATLs significantly increased colocalization of TTR^V30M^ with p62 (Fig. 4H and 4I). However, dose-response immunoblotting analyses showed no significant reduction in TTR^V30M-MYC^ levels at ATL concentrations up to 2.5 μM (Fig. 4J and 4K). These findings suggest that although chemical activation of p62 facilitates the sequestration of amyloidogenic TTR and limits further accumulation, it is insufficient to induce efficient degradation of TTR aggregates.

## Targeted protein degradation of both intracellular and transmitted extracellular amyloidogenic mutant TTR using Autotacs

Because R-BiP acts as a molecular bridge between p62 and TTR, we next sought to pharmacologically mimic and accelerate this process by developing a heterobifunctional chimeric degrader for hATTR using the AUTOTAC platform [56]. As a first step to identify an appropriate target-binding ligand (TBL) for mutant TTR, we generated a set of Autotacs carrying different TBLs. Three compounds were selected based on predicted binding modes of their TBLs to TTR or misfolded protein regions (Fig. 5A). ATC201 employs curcumin, which selectively binds the T4 pocket of TTR, stabilizing its native conformation and preventing oligomer formation. ATC202 adopts a de-brominated derivative of the oligomeric modulator anle138b, which selectively occupies beta-beta sheet signatures in disordered aggregates [60]. ATC203 uses 4-phenylbutyric acid, which binds exposed hydrophobic patches common to misfolded proteins [61]. Among these, molecular mechanics/generalized born surface area (MM-GBSA) analysis with mutant TTR showed that ATC201 exhibited the most favorable binding affinity (MM-GBSA score: –62.11 kcal/mol), superior to that of the TBL curcumin alone (Table S1).

**Figure 5.**
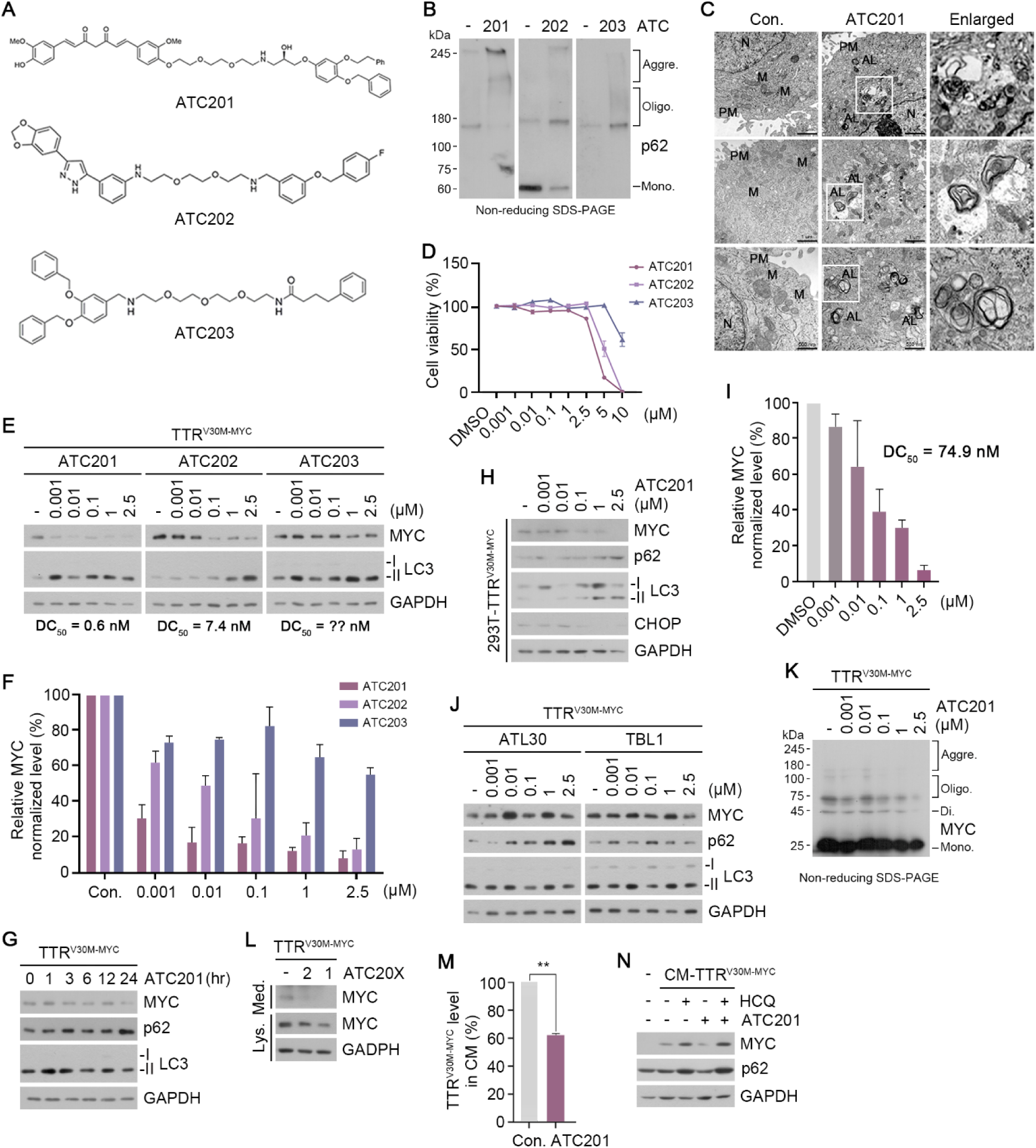
Targeted degradation of extra-/intracellular deposited TTR *via* AUTOTAC. (**A**) Chemical structures of ATC201, ATC202 and ATC203. (**B**) *In vitro* oligomerization assay in HEK293T cells incubated with the indicated compounds (1 μM, 24 h). (**C**) TEM of HEK293T cells treated with ATC201 (1 μM, 24 h). Scale bar represents 1 μm (first and second rows), 500 nm (third row). (**D**) Cell viability assay in HeLa cells treated with the indicated compounds at the indicated concentrations (24 h) (n=3). (**E**) Immunoblotting analysis of HeLa cells transfected with TTR^V30M-MYC^, subsequently treated with the indicated compounds at the indicated concentrations (24 h). (**F**) Densitometry of TTR levels in (E) (n = 3). (**G**) Immunoblotting analysis of HeLa cells transfected with TTR^V30M-MYC^, subsequently treated with ATC201 at the indicated time points (100 nM). (**H**) Immunoblotting analysis of TTR^V30M-MYC^ expressing 293T stable cell line treated with ATC201 at the indicated concentrations (24 h). (**I**) Densitometry of (H) (n = 3). (**J**) Immunoblotting analysis of HeLa cells transfected with TTR^V30M-MYC^, subsequently treated with ATL30 or TBL1 at the indicated concentrations (24 h). (**K**) *In vivo* oligomerization assay of HeLa cells transfected with TTR^V30M-MYC^, subsequently treated with ATC201 at the indicated concentrations (24 h). (**L**) Immunoblotting analysis of lysates and TCA-precipitated media from HeLa cells transfected with TTR^V30M-MYC^, subsequently treated with ATC201 or ATC202 (1 μM, 24 h). (**M**) ELISA assay of concentrated media from HeLa cells transfected with TTR^V30M-MYC^, subsequently treated with ATC201 (1 μM, 24 h) (n=3). (**N**) Immunoblotting analysis of HeLa cells co-treated with CM containing secreted TTR^V30M-MYC^ and ATC201 (1 μM, 24 h) in the presence or absence of HCQ (10 µM, 24 h). Data are presented as means ± SD. Each n represents an independent biological replicate. Unpaired two-tailed Student’s t test; p* < 0.05, **p < 0.01, ***p < 0.001, ****p < 0.0001 and ns, not significant.

All three Autotacs efficiently induced p62 self-oligomerization *in vitro* (Fig. 5B), confirming successful engagement to p62. Consistent with these results, transmission electron microscopy of ATC201-treated HEK293T cells revealed substantial accumulation of autophagosomes and autolysosomes (Fig. 5C). We next assessed the degradative efficacy of Autotacs using dose-dependent degradation assays in cells expressing TTR^V30M-MYC^. Cell viability assays showed no cytotoxicity for any compound up to 2.5 μM, which was subsequently used as the maximal concentration (Fig. 5D). In HeLa cells transiently expressing TTR^V30M-MYC^, ATC203 showed no detectible degradative efficacy, whereas ATC202 and ATC201 induced robust, concentration-dependent degradation (Fig. 5E and 5F). ATC201 exhibited the highest potency, with a DC_50_ of 0.6 nM and a D_max_ of 10 nM (Fig. 5E and 5F), and reduced TTR^V30M-MYC^ levels by ∼50% within 9 hours at 100 nM (Fig. 5G).

Comparable results were observed in HEK293T cells stably expressing TTR^V30M-MYC^ (DC₅₀, 74.9 nM; Fig. 5H and 5I), as well as in HepG2 cells expressing endogenous TTR and ectopic TTR^V30M-MYC^ (Fig. S5A). Notably, ATC201 treatment also reduced CHOP expression (Fig. 5H), demonstrating its activity to alleviate ER stress under amyloidogenic conditions. This degradative activity was not reproduced by either the ATL or TBL moiety alone (Fig. 5J), indicating that the bifunctional structure of ATC201 is essential for TTR degradation. Under non-reducing SDS-PAGE conditions where disulfide-linked oligomers are preserved, both ATC201 and ATC202 significantly reduced high-molecular-weight TTR^V30M-MYC^ species in a concentration-dependent manner (Fig. 5K, S5B and S5C). These results demonstrate that ATC201 and ATC202 selectively promote degradation of oligomeric and aggregated of TTR^V30M-MYC^ species, thereby mitigating amyloid-driven ER stress.

We also determined whether Autotacs can reduce extracellular TTR in HeLa cells transiently expressing TTR^V30M-MYC^. ATC201 and, to a less extent, ATC202 decreased extracellular TTR levels (Fig. 5L). Consistently, ELISA of CM showed that ATC201 reduced TTR^V30M-MYC^ levels by ∼40% (Fig. 5M). To determine whether this reduction reflects enhanced uptake and degradation, cells were incubated with CM containing TTR^V30M-MYC^ in the presence of ATC201. Under these conditions, internalized extracellular TTR underwent accelerated degradation, which was effectively blocked by HCQ, indicating lysosome dependent clearance (Fig. 5N). These results demonstrate that Autotacs can induce lysosomal degradation of both intra- and extracellular TTR variants.

## ATC201 engages p62 to promote lysosomal clearance of amyloidogenic TTR

To determine whether ATC201 promotes TTR clearance through p62-dependent autophagy, we first evaluated the binding mode *in silico*. Molecular docking and MM-GBSA analyses showed that ATC201 formed a stable ternary complex by simultaneously engaging mutant TTR and the p62 ZZ domain (Fig. 6A and 6B). The curcumin moiety occupied the T4 binding pocket of TTR via hydrogen bonding with Lys15 and Arg104, while the ATL interacted with Val135 of the p62 ZZ domain. The complex exhibited a binding energy of – 71.26 kcal/mol (Supplementary Table 1), indicating high stability and supporting the structural basis for its bifunctional mechanism. In cellular assays, siRNA-mediated depletion of p62 completely abolished ATC201-induced degradation of TTR^V30M–MYC^ (Fig. 6C), whereas knockdown of the Ub-encoding *Ubb* gene had no effect (Fig. 6D), supporting a p62-dependent but Ub-independent mechanism of action. Immunocytochemistry further showed that ATC201 induced formation of p62-TTR puncta (Fig. 6E and 6F), the majority of which were positive for LC3^+^ autophagic membranes (Fig. 6G and 6H). This effect was not observed with either the TBL or ATL moiety alone (Fig. 6G, 6H and S6A), confirming that the bifunctional Autotac structure is required for selective targeting of amyloidogenic TTR to autophagic compartments.

**Figure 6.**
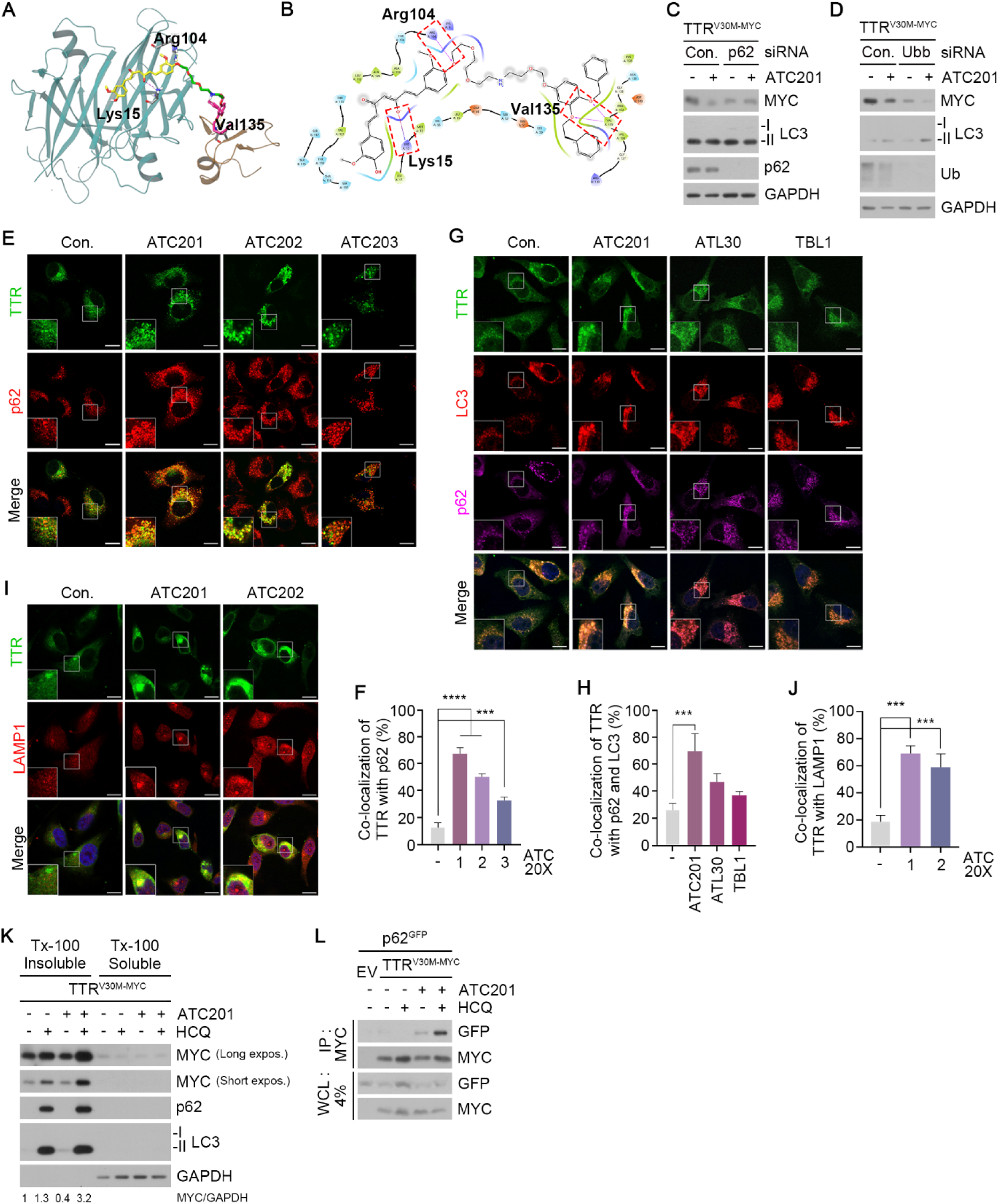
p62-dependent autophagic delivery of TTR. (**A**) Illustration of the detailed 3D docking pose of ATC201 within the mutant TTR-p62 ZZ domain complex. The mutant TTR structure is shown in cyan, while the p62 ZZ domain is depicted in brown. ATC201 features three distinct regions: the curcumin moiety (yellow), a flexible linker (green), and the terminal portion interacting with the p62 ZZ domain (purple). (**B**) Illustration of the 2D interaction diagram of ATC201 with the mutant TTR (Chain A) and the p62 ZZ domain (Chain Z). Key interactions are highlighted in red dashed boxes. (**C-D**) Immunoblotting analysis of HeLa cells transfected with TTR^V30M-MYC^, subsequently treated with ATC201 (1 μM, 24 h) under siRNA-mediated knockdown of *p62* and *Ubb* (40 nM, 48 h). (**E**) Immunostaining analysis of HeLa cells transfected with TTR^V30M-MYC^, subsequently treated with the indicated compounds (1 μM, 24 h). (**F**) Quantification of (E) (n = 50 cells). (**G**) Immunostaining analysis of HeLa cells transfected with TTR^V30M-MYC^, subsequently treated with the indicated compounds (1 μM, 24 h). (**H**) Quantification of (G) (n = 50 cells). (**I**) Immunostaining analysis of HeLa cells transfected with TTR^V30M-MYC^, subsequently treated with ATC201 or ATC202 (1 μM, 24 h). (**J**) Quantification of (I) (n = 50 cells). (**K**) Triton X-100-fractionation assay of HeLa cells transfected with TTR^V30M-MYC^, subsequently treated with ATC201 (1 μM, 24 h) in the presence or absence of HCQ (10 μM, 24 h). (**L**) CoIP assay in HEK293T cells co-transfected with p62^GFP^ and TTR^V30M-MYC^, subsequently treated with ATC201 (1 μM, 24 h) in the presence or absence of HCQ (10 μM, 24h). Scale bar represents 10μm. Data are presented as means ± SD. Each n represents an independent biological replicate. One-way ANOVA; p* < 0.05, **p < 0.01, ***p < 0.001, ****p < 0.0001 and ns, not significant.

We next examined whether Autotacs promote lysosome-dependent clearance of aggregated TTR. When visualized using immunostaining analyses, ATC201 and ATC202 efficiently targeted TTR to LAMP1^+^ lysosomes (Fig. 6I and 6J), suggesting that p62–TTR complexes are directed to autophagic vesicles and subsequently trafficked to lysosomes.

Consistently, ATC201-driven clearance of oligomeric TTR was markedly blocked by hydroxychloroquine, indicating lysosome-dependent degradation (Fig. S6B). When TTR^V30M-MYC^ was fractionated into insoluble or soluble species using Triton X-100, ATC201 preferentially promoted autophagic degradation of insoluble species as compared with soluble species (Fig. 6K). These results demonstrate Autotacs selectively promote the lysosomal degradation of oligomeric and aggregated TTR. To further confirm that ATC201 facilitates lysosomal clearance of TTR via p62 interaction, we performed co-IP under lysosomal blockade. ATC201 enhanced the formation and accumulation of TTR–p62 complexes in the presence of HCQ (Fig. 6L), indicating active autophagic flux. These findings show that Autotacs directs aggregated TTR to lysosomal degradation through a p62-dependent autophagy pathway.

## ATC201 induces the degradation of TTR^V30M^ in hATTR^V30M^ model mice

To assess the physicochemical properties of ATC201 as a therapeutic agent for hATTR, we measured its pharmacokinetic (PK) profiles in ICR mice. Following intravenous injection, ATC201 reached a AUC_last_ of 1,607.2 h*ng/ml and C_max_ of 5,602 ng/ml, with a terminal half-life (T_1/2_) of 3 hours (Supplementary Table 2). Intraperitoneal administration resulted in a C_max_ of 2,092.4 ng/ml and an AUC_last_ of 2,881.7 h*ng/ml (Supplementary Table 2), indicating favorable systemic exposure via clinically relevant routes.

We next evaluated the degradative efficacy of ATC201 in a C57BL/6 transgenic mouse model expressing human TTR^V30M^ [62]. Twelve-month-old mice, corresponding to mid-late hATTR stages, received intraperitoneal injection of 10 mg/kg ATC201 once per week for 3 months (Fig. 7A). Immunoblotting analyses showed that 12 doses of ATC201 reduced TTR^V30M^ levels in the colon, small intestine, liver, brain, and muscle. Among these, TTR^V30M^ levels were most prominently lowered by ∼70% in the colon (Fig. 7B and 7D) and small intestine (Fig. 7C and 7E), the major gastrointestinal target organs in TTR^V30M^-associated FAP [63]. The lowering correlated with elevated levels of p62 (Fig. 7B, 7C, 7F and 7G) and lipidated LC3 (Fig. 7B, 7C, 7H and 7I), consistent with enhanced autophagic activity. In addition, significant reductions (∼40%) in TTR levels were observed in the liver (Fig. S7A and S7B), brain (Fig. S7C and S7D) and muscle (Fig. S7E and S7F), whereas no noticeable efficacy was seen in the lung (Fig. S7G and S7H). Immunohistochemistry (IHC) of the colon revealed that ATC201 markedly reduced both intracellular and extracellular TTR (Fig. 7J and 7K).

**Figure 7.**
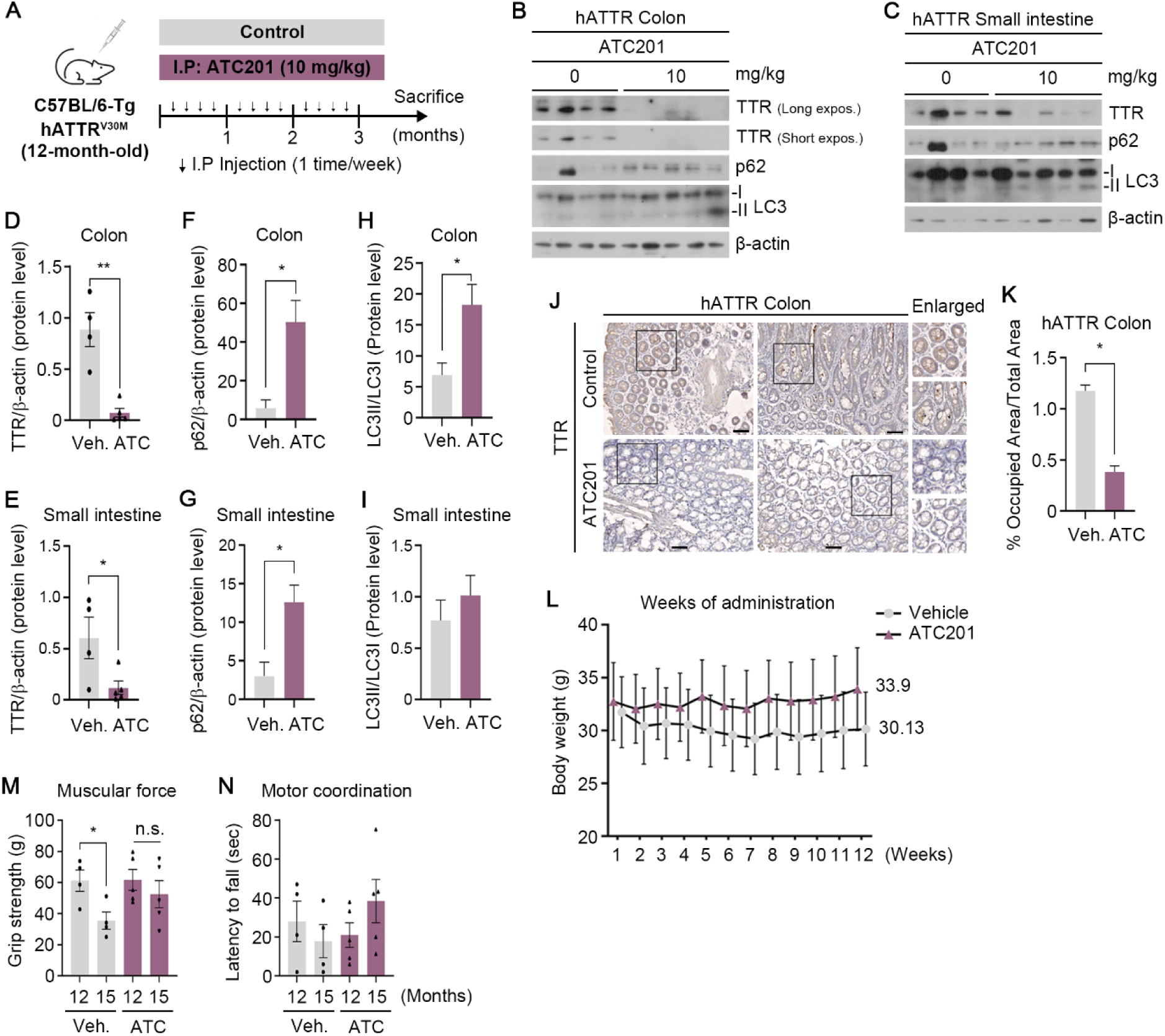
AUTOTAC mitigates amyloidosis in hATTR mouse model. (**A**) Schematic of hATTR^V30M^ murine model with injection timeline and details of ATC201. (**B-C**) Immunoblotting analysis of colon and small intestine tissues from hATTR^V30M^ mice injected with ATC201, as described in (A). (**D-I**) Densitometry of TTR, p62 or LC3II/I levels in (B) and (C) (n = 9). (**J**) Immunohistochemistry analysis of colon tissues from hATTR mice injected with ATC201, as described in (A). (**K**) Quantification of (J). (**L**) Body weight changes during compound administration (Vehicle, n=4; ATC201, n=5). (**M**) Grip strength test examining fore- and hindlimb muscle force (Vehicle, n=4; ATC201, n=5). (**N**) Rotarod test examining motor coordination and balance (Vehicle, n=4; ATC201, n=5). Scale bar represents 50 μm. Data are presented as means ± S.E.M. Each n represents an independent biological replicate. Unpaired two-tailed Student’s t test; p* < 0.05, **p < 0.01, ***p < 0.001, ****p < 0.0001 and ns, not significant.

To further validate the preventive efficacy of ATC201, we employed HM30 transgenic mice, in which mouse TTR was knocked out and human TTR^V30M^ was ectopically overexpressed [64, 65]. Mice aged 7-8 months received intraperitoneal injections with 10 mg/kg ATC201 three times per week for one month (Fig. S8A). Immunoblotting showed that TTR^V30M^ levels were significantly lowered in both the colon (Fig. S8B and S8D) and stomach (Fig. S8C and S8E) following 12 doses. Consistently, IHC revealed significantly weaker TTR staining in ATC201-treated mice compared with controls (Fig. S8, F to I). These results demonstrate the efficacy of ATC201 in degrading TTR aggregates in hATTR^V30M^ model mice.

### ATC201 mitigates neuropathic and weight loss symptoms in hATTR^V30M^ model mice

FAP is characterized by progressive axonal degeneration, neuromuscular dysfunction, and systemic manifestations such as weight loss, primarily driven by the accumulation of TTR aggregates in the peripheral nerves [66]. To assess the therapeutic efficacy of ATC201 *in vivo*, we examined systemic homeostasis and neuromuscular performance in C57BL/6 transgenic mice expressing human TTR^V30M^ treated once weekly for 3 months from 12 to 15 months of age. Following the 3-month regimen, ATC201-treated animals exhibited a modest increase in body weight (+3.4 ± 1.3%), whereas vehicle controls showed a 5.0 ± 4.5% decrease, resulting in an overall ∼8% difference between groups (Fig. 7L). Forelimb grip strength was largely preserved in ATC201-treated mice but declined by ∼42% in vehicle-treated mice (p = 0.018; Fig. 7M). Consistently, ATC201 also markedly improved motor coordination in the rotarod test (+162.0 ± 114.1%) compared with progressive impairment observed in controls (–21.5 ± 22.6%), although the effect did not reach statistical significance (p = 0.185; Fig. 7N). These results suggest that ATC201 preserves systemic and neuromuscular function, supporting its therapeutic potential in hATTR through selective aggregates clearance.

## Discussion

TTR amyloidosis is a progressive systemic disorder characterized by the accumulation of misfolded TTR as extracellular aggregates that spread through systemic circulation and drive multi-organ dysfunctions [4, 67]. Although current therapies such as tetramer stabilizers and RNA interference can delay the *neo*-synthesis of pathogenic TTR monomers, pre-existing aggregates, which drive the aggravating cycle of proteotoxicity, remain beyond their reach [20, 21, 68].

This study was designed to develop a therapeutic strategy targeting extracellular deposition of pathogenic TTR aggregates in hATTR. We show that intracellular TTR^V30M^ undergoes rapid misfolding, secretion, aggregation, and propagation to neighboring cells within 24 hours. The N-degron pathway induces the lysosomal degradation of these misfolded aggregates. In this mechanism, ER-derived BiP undergoes Nt-arginylation in response to incoming cargoes. As an N-degron, the resulting Nt-Arg of R-BiP binds the ZZ domain of p62, allosterically inducing p62 self-polymerization and formation of TTR-p62 condensates destined for lysosomal degradation. Leveraging this mechanism, we applied the AUTOTAC platform to develop a chimeric molecule that bridges TTR aggregates to p62.

ATC201 induced degradation of pathogenic TTR aggregates in hATTR mouse models and improved muscle strength and motor performance. Our results highlight the N-degron pathway as a promising therapeutic target in TTR amyloidosis and potentially other proteinopathies characterized by prion-like propagation of misfolded aggregates.

TTR aggregates are primarily deposited as extracellular amyloid fibrils, where they act as prion-like seeds that promote further aggregation and disseminate systemically across multiple organs [69, 70]. Here, we provide direct evidence that pathogenic TTR undergoes a rapid secretion-transmission cycle. Non-reducing SDS-PAGE revealed that a substantial fraction (approximately 40-60%) of TTR variants secreted within 24 hours re-entered cells, thereby accelerating ER stress and cytotoxicity. The precise molecular route of re-entry in our system remains unresolved. Previous work has shown that ERdj3, an ER-resident Hsp40 co-chaperone, selectively binds misfolding-prone TTR^A25T^, is co-secreted into the extracellular environment, and can promote their internalization [71]. In addition, BiP, calreticulin, and calnexin relocalize to the cell surface or extracellular space, where they engage misfolded proteins [72–74]. These observations raise the possibility that BiP, like ERdj3, may facilitate both internalization and degradation of extracellular TTR aggregates. Although our current data do not yet conclusively establish such a role, they point to an intriguing hypothesis for future studies. Crucially, while much of our mechanistic analysis relied on ectopic TTR^V30M^ expression rather than exogenous uptake, the demonstrated secretion–re-entry cycle supports this framework for investigating post-entry fate.

A major mechanistic insight of this study is the identification of R-BiP as a molecular bridge linking pathogenic TTR to p62 for autophagic degradation. BiP was upregulated following exposure to TTR^V30M^ and physically associated with TTR^V30M^ in an ATPase-dependent manner. BiP overexpression and consequently increased BiP-TTR interaction reduced both intracellular and extracellular TTR. Subsequent Nt-arginylation of BiP by ATE1 enabled its engagement with the p62 ZZ domain, thereby coupling misfolded TTR to autophagic vesicles. Disruption of BiP ATPase activity, ATE1 function, or the p62 ZZ domain abolished this clearance route, highlighting R-BiP as a key molecular bridge between pathogenic TTR and the autophagy machinery. Notably, each TTR mutant exhibited distinct aggregation-prone behaviors, ranging from monomers and oligomers to aggregates. The existence of more than one hundred mutant conformations unique to TTR begs the question of how cells selectively recognize and metabolize mutant TTR according to their conformation. One possibility is that since multiple ER chaperones such as calreticulin, protein disulfide isomerase and others are known to be retrotranslocated to the cytosol and Nt-arginylated by ATE1 [29], the scope of the N-degron pathway in combating TTR amyloidosis may be expanded to a wide myriad of TTR mutants beyond what were used in this study. Our findings, thus, reveal new insights into the role of quality control pathways in regulating secretion and extracellular aggregation of amyloidogenic TTRs, and may pave the way for alternative therapeutic strategies that pharmacologically modulate ER chaperones.

While TTR expression and accumulation have been associated with increased levels of autophagy-related factors including p62, lipidated LC3 and BECN1 [75], TTR aggregates are known to impair autophagy [76]. We speculated that autophagy is initially activated in response to TTR accumulation as a cytoprotective mechanism. Indeed, TTR overexpression facilitated autophagy induction and accelerated autophagic flux, as indicated by increased p62 and lipidated LC3 levels. Pharmacological activation of p62 by N-degrons promoted p62 oligomerization and colocalization with TTR but failed to substantially reduce TTR aggregates. In contrast, ATC201, composed of a curcumin-based TTR-binding ligand and a chemical N-degron, not only enhanced autophagic activity but also facilitated degradation of intracellular TTR. ATC201 also reduced extracellular TTR by ∼40%, accelerating lysosomal clearance of re-entered aggregates. This bifunctional activity was not recapitulated by either TBL or ATL alone. These results indicate that simply stimulating autophagy is insufficient to eliminate TTR aggregates due to limited cargo selectivity, and that both target-binding and autophagy-inducing activities are required for effective targeted protein degradation TTR pathology.

Despite the compelling efficacy of ATC201 in degrading TTR aggregates and mitigating disease phenotypes, several considerations remain. First, much of our mechanistic work relied on ectopically expressed TTR^V30M^ rather than CM-derived TTR. While CM-based assays confirmed intercellular transmission, technical challenges associated with CM use in all experiments led us to focus primarily on overexpression systems. This approach allowed reproducibility and mechanistic dissection but may not fully recapitulate the dynamics of secreted TTR aggregates *in vivo*. Second, tissue-specific variability, including minimal efficacy in the lung, suggests that local pharmacokinetics and autophagic capacity shape therapeutic outcomes. Functional improvements were broadly concordant but not uniformly significant, indicating that larger, longer-term studies will be needed to confirm benefit. Finally, chronic modulation of p62-dependent autophagy requires careful safety evaluation given its central role in proteostasis. Nonetheless, by directly linking the fate of transmitted TTR to the N-degron pathway and demonstrating its pharmacological activation via AUTOTAC, this study provides both mechanistic insight and a translational proof-of-concept. The modularity of AUTOTAC further opens the possibility of extending selective aggregate clearance to other amyloidogenic proteins, including tau, α-synuclein, and TDP-43.

From a translational perspective, ATC201 exhibits several properties that support its potential advancement toward clinical application in hATTR. PK analyses in mice demonstrated that ATC201 achieves favorable systemic exposure following both intravenous and intraperitoneal administration, with a T_1/2_ of ∼3 hours and robust C_max_ and AUC values, indicating sufficient bioavailability to engage tissue-deposited TTR aggregates. Notably, a low-frequency dosing regimen (10 mg/kg once weekly for 12 weeks) was sufficient to induce marked reductions in TTR aggregate burden and to ameliorate neuromuscular dysfunction in hATTR mouse models, suggesting that intermittent dosing may be adequate for sustained therapeutic benefit. This dosing paradigm contrasts favorably with existing therapies, such as TTR tetramer stabilizers or RNA interference–based agents, which primarily suppress de novo TTR production but do not eliminate pre-existing amyloid deposits and require lifelong continuous administration. By directly targeting pathogenic aggregates for lysosomal degradation, ATC201 offers a disease-modifying strategy that addresses a major unmet need in hATTR management. To advance ATC201 toward clinical development, several steps will be essential, including optimization of formulation and delivery, comprehensive toxicology and safety pharmacology studies, and evaluation of long-term effects of sustained p62-dependent autophagy activation. In addition, expanded PK and pharmacodynamic (PD) analyses in larger animal models, as well as assessment of combination strategies with current standard-of-care therapies, may further refine its clinical positioning. Collectively, these considerations support the feasibility of translating AUTOTAC-based degraders such as ATC201 into first-in-class autophagy-targeting therapeutics for hATTR and potentially other systemic proteinopathies driven by pathogenic protein aggregation.

## Materials and Methods

### Antibodies

Primary antibodies used were as follows: anti-FLAG_M2 mouse Ab (Sigma-Aldrich, F1804), anti-FLAG rabbit Ab (Sigma-Aldrich, SAB4301135), anti-Myc rabbit Ab (71D10) (Cell Signaling Technology, 2278S), anti-GAPDH rabbit Ab (Bioworld Technology, AP0066), anti-β-actin mouse Ab (Sigma-Aldrich, A1978), anti-C-Myc (9E10) mouse Ab (Santa Cruz Biotechnology, sc-40), anti-hATE1 (H-12) mouse Ab (Santa Cruz Biotechnology, sc-271219), anti-TTR rabbit Ab (Abcam, D527001), anti-p62 mouse Ab (Abcam, ab56416), anti-p62 rabbit Ab (Santa Cruz Biotechnology, sc-25575), anti-BiP rabbit Ab (Abcam, ab21685), anti-CRT mouse Ab (Abcam, ab2907), anti-R-BiP rabbit Ab (Cha-Molstad et al., 2015), anti-WIPI2 rabbit Ab (Cell Signaling Technology, 8567), anti-ATG5 rabbit Ab (Sigma-Aldrich, A0856), anti-LAMP1 rabbit Ab (Sigma-Aldrich, L418), anti-CHOP mouse Ab (Cell Signaling Technology, 2895), and anti-PARP rabbit Ab (Cell Signaling Technology, 9542).

The secondary antibodies used were as follows: anti-rabbit IgG-HRP (Cell Signaling Technology, 7074), anti-mouse IgG-HRP (Cell Signaling Technology, 7076), Alexa Fluor 488 goat anti-rabbit IgG (Invitrogen, A11034), Alexa Fluor 488 goat anti-mouse IgG (Invitrogen, A11029), Alexa Fluor 555 goat anti-rabbit IgG (Invitrogen, A21429), and Alexa Fluor 555 goat anti-mouse IgG (Invitrogen, A32727).

Control antibodies were used: Normal mouse IgG (Santa Cruz Biotechnology, sc-2025), and Normal rabbit IgG (Cell Signaling Technology, 2729S).

### Plasmids, siRNAs and primers

The following plasmids were used: TTR^MYC^ (Origene Technologies, RC204976), pLV-EF1α-IRES-Puro (Addgene, 85132), psPAX2 (Addgene, 12260), pMD2.G (Addgene, 12259), pEGFP-N3/Ub-R-BiP-GFP (This paper), pEGFP-N3/Ub-V-BiP-GFP (This paper), pcDNA3.1-myc-His-B/p62 (Cha-Molstad et al., 2015), and pcDNA3.1-myc-His-B/p62ΔZZ (Cha-Molstad et al., 2015).

The following oligonucleotides and siRNAs were used: siRNA negative control (Bioneer, SN-1003), siRNA against p62 (Bioneer, 1144479), siRNA against *ATE1* (Thermo Fisher Scientific, 4390824), siRNA against *ATG5* (Thermo Fisher Scientific, 4392420), *GAPDH* primer (Cosmogenetech; Fwd: 5′-TCAACAGCGACACCCACTCC-3′, Rvs: 5′-TGAGGTCCACCACCCTGTTG-3′), *CHOP* primer (Cosmogenetech; Fwd: 5′-GGAGCTGGAAGCCTGGTATG-3′, Rvs: 5′-GCAGGGTCAAGAGTGGTGAA-3′), *BiP* primer (Cosmogenetech; Fwd: 5′-AATGACCAGAATCGCCTGAC-3′, Rvs: 5′-CGCTCCTTGAGCTTTTTGTC-3′), *XBP1* primer (Cosmogenetech; Fwd: 5′-AACCAGGAGTTAAGACAGCGCTT-3′, Rvs: 5′-CTGGGTCCAAGTTGTCCAGAAT-3′), and *ATF4* primer (Cosmogenetech; Fwd: 5′-ATGACCGAAATGAGCTTCCTG-3′, Rvs: 5′-GCTGGAGAACCCATGAGGT-3′).

### Cell culture

HepG2 cells (KCLB, 88065) were obtained from Korean Cell Line Bank. HeLa cells (ATCC, CCL-2) and HEK293T cells (ATCC, CRL-11268) were obtained from American Type Culture Collection. Cell lines used in this study were maintained in Dulbecco’s Modified Eagle’s Medium (DMEM; Gibco, 11995065) supplemented with 10% Fetal Bovine Serum (FBS; Gibco, 12483020) and antibiotics (100 U/mL penicillin and 100 μg/mL streptomycin; Thermo Fisher Scientific, 15140122). All cell lines were maintained at 37°C and 5% CO_2_ in an incubator.

### Generation of stable cell line

Lentiviral vectors encoding WT TTR or TTR^V30M^ tagged with Myc were constructed using the pLV-EF1α-IRES-Puro backbone (Addgene, 85132). To produce lentiviral particles, HEK293T cells were co-transfected with the expression constructs and packaging plasmids psPAX2 (Addgene, 12260) and pMD2.G (Addgene, 12259) using Lipofectamine 3000 (Thermo Fisher Scientific, 13778150). Virus-containing supernatants were collected 48 hours post-transfection, filtered through a 0.45 μm filter, and used to transduce HEK293T cells. Transduced cells were selected with 1 μg/ml puromycin (InvivoGen, ant-pr-1) for 3–5 days until uninfected control cells were eliminated. Stable cell lines were verified using immunoblotting and/or immunocytochemistry.

### Transfection of nucleic acids

An appropriate amount of plasmids was transfected into the cells by using Lipofectamine 2000 (Thermo Fisher Scientific, 11668019) and siRNAs were transfected at 40 nM using Lipofectamine RNAiMAX (Thermo Fisher Scientific, 13778030) according to the manufactural instruction. For co-transfection of plasmids and siRNAs, Lipofectamine 2000 was used. The lipofectamine reagents and nucleic acids were incubated in Opti-MEM (Gibco, 31984-070) respectively for 5 min, followed by mixing and incubating for 15 min. 5 hours after treatment of the transfection mixture, transfected cells were washed with fresh media and incubated for a desired time.

### Protein degradation cycloheximide-chase assay

For pulse chase analysis, cells at 80% confluence were transfected or treated with indicated plasmids and reagent, followed by a chase in the presence of cycloheximide (CHX; 10 μg/ml; Sigma-Aldrich, 01810) at the indicated time points.

### Conditioned media collection and treatment

Conditioned media (CM) was collected from HeLa or HEK293T cells that were transfected with TTR plasmids. After 24 hours of transfection, the medium was replaced with fresh media. The medium was collected after 24 hours and centrifuged at 100 g to remove cell debris and dead cells. Supernatants were collected and subsequently treated to fresh HeLa or HEK293T cells to assess the extent of TTR propagation to these new cells.

### Trichloroacetic acid precipitation of media

Media were harvested from transfected HeLa cells centrifuged at 2,000 g for 10 min at 4 °C to remove cell debris. The clarified supernatant was transferred to fresh tubes, and proteins were precipitated by the addition of ice-cold 100% trichloroacetic acid (TCA; Biosaesang, T1085) to a final concentration of 10% (v/v).

Samples were incubated on ice for 30 min and centrifuged at 14,000 g for 15 min at 4 °C. The resulting protein pellets were washed twice with cold acetone, air-dried briefly, and resuspended in 4X Laemmli sample buffer (Bio-Rad Laboratories, 1610747), followed by boiling 5 min at 100°C, and separated on SDS–PAGE for immunoblotting analysis.

### Preparation of purified protein

HEK293T cells were transfected with a plasmid encoding WT TTR or TTR^V30M^ tagged with Myc using Lipofectamine 3000 in T175 flasks, according to the manufacturer’s instructions. 48 hours after transfection, culture media were harvested, centrifuged at 2,000 g for 10 min to remove cell debris, and filtered through a 0.45 μm syringe filter. The supernatant was incubated with anti-MYC agarose beads (Thermo Fisher Scientific, 20168) for 2–4 hours at 4°C with gentle rotation. After extensive washing with PBS, bound TTR^MYC^ was eluted using 0.1 M glycine-HCl (pH 2.5) and immediately neutralized with 1 M Tris-HCl (pH 8.0). To remove pre-formed aggregates and enrich soluble monomeric species, the eluted protein was diluted in PBS and subjected to ultracentrifugation at 100,000 g for 1 hour at 4 °C using a TLA-100.3 rotor (Beckman Coulter, SCR_018671).

The resulting supernatant was carefully collected, quantified by Pierce BCA Protein Assay Kit (Thermo Fisher Scientific, 23227), and directly used for downstream applications.

### Cell lysis for total protein

Adherent cells were washed 3 times with cold PBS and lysed by using RIPA buffer (Biosaesang, RC2002-050-00) supplemented with protease inhibitor cocktail (Abbkine Scientific, BMP1001) and phosphatase inhibitor cocktail (Sigma-Aldrich, P5726). Lysates were incubated on ice for 30 min with occasional vortexing and clarified by centrifugation at 15,000 g for 15 min at 4 °C. Supernatants were collected and protein concentrations were determined using Pierce BCA Protein Assay Kit, and equal amounts of protein were used for Thioflavin T (ThT) fluorescence assays and immunoblotting. For immunoblotting, 4X Laemmli sample buffer was added to the supernatant, followed by boiling 5 min at 100°C, and separated on SDS–PAGE for immunoblotting analysis.

### Thioflavin T binding assays

Thioflavin T (ThT) binding assays were conducted using either whole-cell lysates or purified recombinant TTR proteins and performed in 96-well black plates using glycine–NaOH buffer (100 mM, pH 8.5). Samples (40 μl per well; final protein concentration: 0.15 mg/ml) were mixed with 50 μl of freshly prepared 10 μM ThT (Sigma-Aldrich, T3516) in the same buffer and incubated for 5 min at room temperature in the dark. For cell-based assays, ThT fluorescence was recorded immediately after incubation. For in vitro assays, purified TTR proteins were incubated at 37 °C, and ThT fluorescence was monitored every 15 min over a 72-hour period to assess amyloid formation kinetics.

Fluorescence was measured using a Tecan plate reader (Tecan, SCR_020299) (excitation 450 nm, emission 490 nm), and PBS was used as a blank control. All measurements were performed in triplicate.

### Triton X-100 soluble/insoluble fractionation

Adherent cells were washed 3 times with cold PBS and lysed using Triton X-100 lysis buffer (20 mM HEPES pH 7.9, 0.2 M KCl, 1 mM MgCl_2_, 1 mM EGTA, 1% Triton X-100 [Biosaesang, TR1020-500-00], 10% glycerol, protease inhibitor cocktail and phosphatase inhibitor cocktail) and incubated on ice for 15 minutes, followed by centrifugation at 15,000 g for 15 min at 4 °C. The supernatant was collected as the soluble fraction and the pellet as the insoluble fraction. The insoluble fraction was washed four times with PBS and then lysed with SDS-detergent lysis buffer (20 mM HEPES pH 7.9, 0.2 M KCl, 1 mM MgCl2, 1 mM EGTA, 1% Triton X-100, 1% SDS, 10% glycerol, protease inhibitor cocktail and phosphatase inhibitor cocktail). Both soluble and insoluble fractions were mixed with 4X Laemmli sample buffer, boiled for 5 min at 100 °C, and separated on SDS–PAGE for immunoblotting analysis.

### In vivo oligomerization

To lyse the cells, the cells were resuspended in lysis buffer (50 mM HEPES pH 7.4, 0.15 M KCl, 0.1% Nonidet P-40, 10% glycerol, protease inhibitor cocktail and phosphatase inhibitor cocktail) and subjected to a cycle of freezing/thawing 10 times.

The cells were centrifuged at 13,000 g for 10 min after 30 min incubation on ice for supernatant collection. Protein concentration was determined using the Pierce BCA Protein Assay Kit. Next, the samples were mixed with non-reducing 4X LDS sample buffer (Thermo Fisher Scientific, 84788), heated, and resolved on a 4-20% gradient SDS-PAGE gel (Bio-Rad, 4561096) for immunoblotting analysis.

### In vitro p62 oligomerization

HEK 293T cells were transfected with a plasmid encoding p62-myc/his fusion proteins [29]. To lyse the cells, the cells were resuspended in lysis buffer (50 mM HEPES pH 7.4, 0.15 M KCl, 0.1% Nonidet P-40, 10% glycerol, protease inhibitor cocktail and phosphatase inhibitor cocktail) and subjected to a cycle of freezing/thawing 10 times. The cells were centrifuged at 13,000 g for 10 min after 30 min incubation on ice for supernatant collection. Protein concentration was determined using the Pierce BCA Protein Assay Kit. Next. A total of 1 µg of protein was incubated with 1 mM p62-ZZ ligands and 100 µM bestatin hydrochloride (Sigma, B8385-5MG) for 2 hours at room temperature. The samples were mixed with non-reducing 4X LDS sample buffer, heated, and resolved on a 4-20% gradient SDS-PAGE gel for immunoblotting analysis.

### Co-immunoprecipitation

HEK293T cells were transfected for exogenous co-IP with indicated constructs using Lipofectamine 2000. After 24 hours, cells were treated with specified reagents for indicated incubation times for both endogenous and exogenous co-IP. The cells were scraped and pelleted by centrifugation, followed by resuspension and lysis with immunoprecipitation buffer (50 mM Tris-HCl pH 7.5, 150 mM NaCl, 0.5% Triton X-100, 1mM EDTA, 1mM phenylmethylsulfonyl fluoride [PMSF; Thermo Fisher Scientific, 36978]), protease inhibitor cocktail and phosphatase inhibitor cocktail for 30 min using rotator at 4 °C. The cells were passed through a 26-gauge 1 mL syringe 10 times and centrifuged at 13,000 g at 4°C. The resulting supernatant were collected and incubated with normal mouse IgG (Santa Cruz Biotechnology, sc-2025) for overnight at 4°C on a rotor and protein A/G-Plus agarose beads (Santa Cruz Biotechnology, sc-2003) were added for 2 hours to preclear the lysate. Cell lysate was then incubated with M2 FLAG-affinity Gel agarose beads (Sigma-Aldrich, A2220) or Myc magnetic beads (Thermo Fisher Scientific, 88842) for 3 hours and the gel beads were washed four times with IP buffer. The proteins bound to beads were dissociated in 4X Laemmli sample buffer, heated at 100 °C for 5 min and separated on SDS–PAGE for immunoblotting analysis.

### Immunoblotting Analysis

A same amount of proteins was loaded on the poly-acrylamide gel and separated by the sodium dodecyl sulfate (SDS)-electrophoresis, followed by transfer onto the polyvinylidene difluoride (PVDF; Millipore, IPVH00010) membranes. For blocking, the membrane was incubated with 4% skim milk in PBST (PBS supplemented with Tween 20 [Duchefa Biochemie, P1362.0500]) solution for 30 min at room temperature, followed by incubation with primary antibodies overnight at 4°C. Subsequently, the membrane was incubated with host specific horseradish peroxidase (HRP)-conjugated secondary antibodies for 1 hour. Protein bands were visualized by using SuperSignal West PICO PLUS (Thermo Fisher Scientific, 34578) and the chemiluminescence signal was detected by using Luminescent image analyzer. Densitometry of bands was analyzed with ImageJ (National Institutes of Health).

### RNA extraction and RT-PCR analysis

Cultured cells were washed with cold PBS 3 times and directly lysed by TRIzol (Thermo Fisher Scientific, 15596026). After incubation for 5 min at RT, chloroform (Sigma-Aldrich, C2432-25ML) was added to the lysate and briefly mixed. The lysates were centrifuged at 12,000 g for 15 min at 4°C. The transparent supernatants were transferred to new tubes and added with the same volume of isopropanol. After mixing vigorously, the mixtures were centrifuged at 12,000 g for 10 min at 4°C. The supernatants were removed leaving white pallets on the bottom of the tubes. After 75% ethanol was added to the pallets, samples were centrifuged at 7,200 g for 5 min. The pallets were dried and dissolved within RNase-free water. The isolated RNA was heated for 10 min at 55°C and quantified using spectrophotometer. 1 μg of RNA were subjected to cDNA synthesis. The mRNA was reverse-transcribed to complementary DNA (cDNA) by using oligo d(T) primers and reverse-transcriptases (SMOBIO Technology, RP1400). The reverse-transcription was performed at 45°C for 50 min and terminated at 85°C for 5 min. The synthesized cDNA was diluted in ultrapure water at 1:7 ratio to analyze by quantitative PCR.

Forward and reverse primers and cDNA were mixed within the qPCR master mix containing SYBR green (SMOBIO Technology, TQ1201). The detailed PCR cycle was followed the user instruction of qPCR kit. Data were analyzed using 2^ΔΔ^ threshold cycle (Ct) method where human *GAPDH* was used for normalization. The primers used in qPCR analyses are listed on key resources table.

#### ELISA

Media were collected from HeLa or HEK293T cells that were transfected with TTR plasmids. After 24 hours of transfection, the medium was replaced with fresh media. The medium was collected after 24 hours and centrifuged at 100 g to remove cell debris and dead cells. The supernatant was concentrated using Amicon 10 K MWCO filters (Merck Millipore, UFC910024). The amounts of TTR in medium were measured by enzyme-linked immunosorbent assay (ELISA) with an ELISA kit (Abcam, ab108895) following the provider’s instruction.

### Immunocytochemistry (ICC)

Cells were cultured on 12 mm coverslips and treated with experimental conditions. At the moment of harvests, the cells were washed with cold PBS 3 times and fixed with 4% paraformaldehyde (PFA; Biosesang, PC2031-050-00) for 15 min at room temperature. After washing three times with PBS, the cells were permeabilized with 0.5% Triton X-100 for 15 min. After 3 times with PBS, the cells were incubated with blocking solution (2% BSA in PBS) for 1 hour and then with primary antibody overnight at 4°C. The cells were washed 3 times for 10 min with PBS and then incubated with secondary antibody conjugated with fluorescent dye for 1 hour. The coverslips were washed 3 times with PBS and mounted on slide glasses using mounting solution containing DAPI (ImmunoBioScience, AR-6501-01). The edge of the coverslips was sealed with the transparent enamel. Confocal images were taken using an LSM 980 airyscan2 (Carl Zeiss, SCR_023915) at 63× magnification with the 488 nm laser, 555 nm laser, or the 405 nm laser and analyzed with the Zeiss ZEN microscope software (Blue edition, SCR_013672). The level of intensity colocalization was assessed using the colocalization tool within the software.

### Transmission electron microscopy (TEM)

The cells were scraped from the culture dish, pelleted via centrifugation, and then resuspended in 2.5% glutaraldehyde in 0.1M sodium cacodylate buffer (pH 7.4) from Electron Microscopy Sciences, and left overnight at 4°C for fixation. Ultra-thin section of the pellets was prepared with ultramicrotome (Leica Microsystems, SCR_020611). Images were captured on TEM electron microscope JEM-1400 at Electron Microscopy Center, Seoul National University Hospital Biomedical Research Institute.

### Cell viability assay

5×10^4^ cells/well were seeded in a 96-well plate and incubated for 1 day and treated with experimental conditions. The cells were incubated with D-Plus™ CCK cell viability assay kit (Dongin Biotech, CCK-3000) for 2 hours in a 37°C incubator. The optical density (OD) value of samples was measured at 450 nm.

#### Molecular mechanics/generalized born surface area (MM-GBSA) analysis

##### Protein structure preparation and mutagenesis

The crystal structure of human TTR bound to ferulic acid and curcumin (PDB ID: 4PME) was obtained from the Protein Data Bank. To generate the amyloidogenic variant, a Val30Met (V30M) mutation was introduced using the Protein Preparation Wizard in Schrödinger Maestro (v14.2). Hydrogen atoms were added, missing side chains were reconstructed, protonation states were adjusted to pH 7.4, and hydrogen bond networks were optimized. The structure of the p62 ZZ domain (PDB ID: 7R1O) was prepared using the same protocol.

##### Ligand preparation and energy minimization

The structure of the bifunctional small molecule Autotacs was generated from its SMILES representation using Schrödinger LigPrep. Protonation states were assigned at physiological pH (7.4), and energy minimization was carried out using the OPLS_2005 force field, with convergence criteria set to an RMSD of 0.01 Å.

##### Receptor grid generation and molecular docking

A receptor grid was generated at the protein–protein interface of the TTR^V30M^–p62 ZZ complex. Molecular docking of Autotacs was performed using Schrödinger Glide in standard precision (SP) mode, allowing ligand flexibility while keeping the receptor rigid. The OPLS_2005 force field was applied throughout the docking procedure.

##### Binding free energy calculations

To evaluate the binding affinity of Autotacs to the TTR^V30M^–p62 ZZ complex, MM-GBSA calculations were conducted using the Prime module in Schrödinger. The ligand–protein complexes were minimized prior to energy decomposition, and total binding energies were computed by summing contributions from van der Waals, electrostatic, solvation, and surface area energies. Protein–ligand interaction diagrams were generated using the Maestro visualization suite.

### Animal

C57BL/6-Tg (TTR^V30M^)14 Imeg frozen embryos were purchased from the Center for Animal Resources and Development (CARD) and transferred into pseudopregnant females by oviductal injection. Genomic DNA from tail biopsies was amplified using human TTR^V30M^-specific primers to confirm transgene integration. For therapeutic assessment, 12-month-old transgenic mice were randomly assigned to vehicle or treatment groups and administered ATC201 (10 mg/kg, intraperitoneally) once per week for 12 weeks.

To assess the preventive efficacy of ATC201, 7–8-month-old HM30-Tg (TTR^V30M^) mice, which lack endogenous mouse TTR and express only human TTR^V30M^, were treated with ATC201 (10 mg/kg, intraperitoneally) three times per week for 4 weeks.

All mice were maintained under specific pathogen-free (SPF) conditions in a temperature-and humidity-controlled facility with a 12-h light/dark cycle and ad libitum access to food and water. Experimental procedures using C57BL/6-Tg (TTR^V30M^) 14Imeg mice were approved by the Institutional Animal Care and Use Committee of Seoul National University (SNU-200812-1). Experiments involving HM30 mice were conducted under authorization 14982/2017 from the Portuguese General Veterinarian Board and were compliant with Directive 2010/63/EU on animal welfare.

### Immunohistochemistry (IHC)

Paraffin-embedded sections of stomach and colon were deparaffinized in xylene (Duksan Pure Chemicals, X0019) and rehydrated through a graded ethanol series. Antigen retrieval was performed in citrate buffer (pH 6.8) at 95 °C for 15 min, followed by quenching of endogenous peroxidase activity using 3% hydrogen peroxide (Millipore, 88597) in methanol. Tissue sections were blocked in PBS containing 10% FBS, 1% BSA, and 0.5% Triton X-100. Sections were incubated overnight at 4 °C with either rabbit monoclonal anti-human TTR antibody or rabbit polyclonal anti-human TTR antibody, followed by HRP-conjugated secondary antibodies for 1 hour at room temperature. Signal was developed using 3,3′-diaminobenzidine (DAB substrate kit, Vector Laboratories, SK-4100), and nuclei were counterstained with hematoxylin (Merck, MHS32). After staining, slides were dehydrated through an ethanol gradient (50%, 70%, 85%, 95% for 5 min each, and 100% twice for 5 min), cleared in xylene (20 min × 2), and mounted with Entellan® (Merck, 107960). Images were acquired using ZEISS® Axio Scan.Z1 (SCR_019139) or Olympus BX50 microscope (SCR_018949) and analyzed with Image Pro Plus software (Media Cybernetics, SCR_007369). The DAB-positive area was quantified and normalized to total image area; results are presented as mean ± SEM.

### Behavior test

All behavioral tests were performed in accordance with protocols approved by the Institutional Animal Care and Use Committee (IACUC). Experiments were conducted using the C57BL/6-Tg (TTR^V30M^)14Imeg mouse model. Forelimb muscle strength was assessed using a BIO-GS3 grip strength meter (Bioseb, SCR_017198), with each mouse subjected to three consecutive trials; the average value was used for analysis. Motor coordination was evaluated using a BX-ROD accelerating rotarod apparatus (Bioseb, SCR_015921), with the rotation speed increasing from 4 to 40 rpm over 300 s. Each mouse underwent three trials per session, and the best performance was recorded.

### Statistics analysis

The data are presented as the mean ± SD or S.E.M. of at least three independent experiments. For each experiment, sample size (n) was determined as stated in the figure legends. Statistical significance was determined using two-tailed unpaired Student’s *t*-test, or one-way ANOVA followed by Tukey’s post hoc test with Prism 9 software (Graph Pad, SCR_002798). P-values were considered significant as follows: ns, not significant; *p<0.05, **p<0.01, ***p<0.001, and ****p<0.0001.

## Acknowledgments

We would like to thank the Y.T.K. and laboratories’ members and AUTOTAC Bio Inc.’s employees for their comments during this study.

## Author contributions

HYK and CHJ designed the study. HYK conducted most of the experimental work. HYK, DYP and YSS contributed to technical assistance in experimental work. SHK and KWS contributed to animal study. HSM, MJS, and MRA provided animal data. HYK wrote this paper. YTK and CHJ provided assistance in review and editing this paper.

## Data availability statement

All data generated or analyzed during this study are included in this article and its Supplementary Information files. Additional datasets that are not shown are available from the corresponding author (Yong Tae Kwon) upon reasonable request.

## Abbreviations

ATE1: Arginyl-tRNA–protein transferase 1; ATC201: Autotac 201; ATC202: Autotac 202; ATL: Autophagy-targeting ligand; ATTR: Transthyretin amyloidosis; AUTOTAC: AUTOphagy-TArgeting Chimera; CM: Conditioned media; co-IP: co-immunoprecipitation; ER: Endoplasmic reticulum; FAP: Familial amyloid polyneuropathy; GFP: Green fluorescent protein; hATTR: Hereditary transthyretin amyloidosis; HCQ: Hydroxychloroquine; HSPA5/BiP/GRP78: Heat shock protein family A (Hsp70) member 5; IHC: Immunohistochemistry; LC3: Microtubule-associated protein 1 light chain 3; MM-GBSA: Molecular Mechanics/Generalized Born Surface Area; Nt: N-terminal; SQSTM1/p62: sequestosome 1; PROTAC: proteolysis-targeting chimera; R-BiP: N-terminally arginylated BiP; T4: Thyroxine; TBL: Target-binding ligand; TPD: Targeted protein degradation; TTR: Transthyretin; Ub: Ubiquitin; UPS: Ubiquitin-proteasome system; ZZ: ZZ-type zinc finger

## Supplementary figures

**Figure S1.**
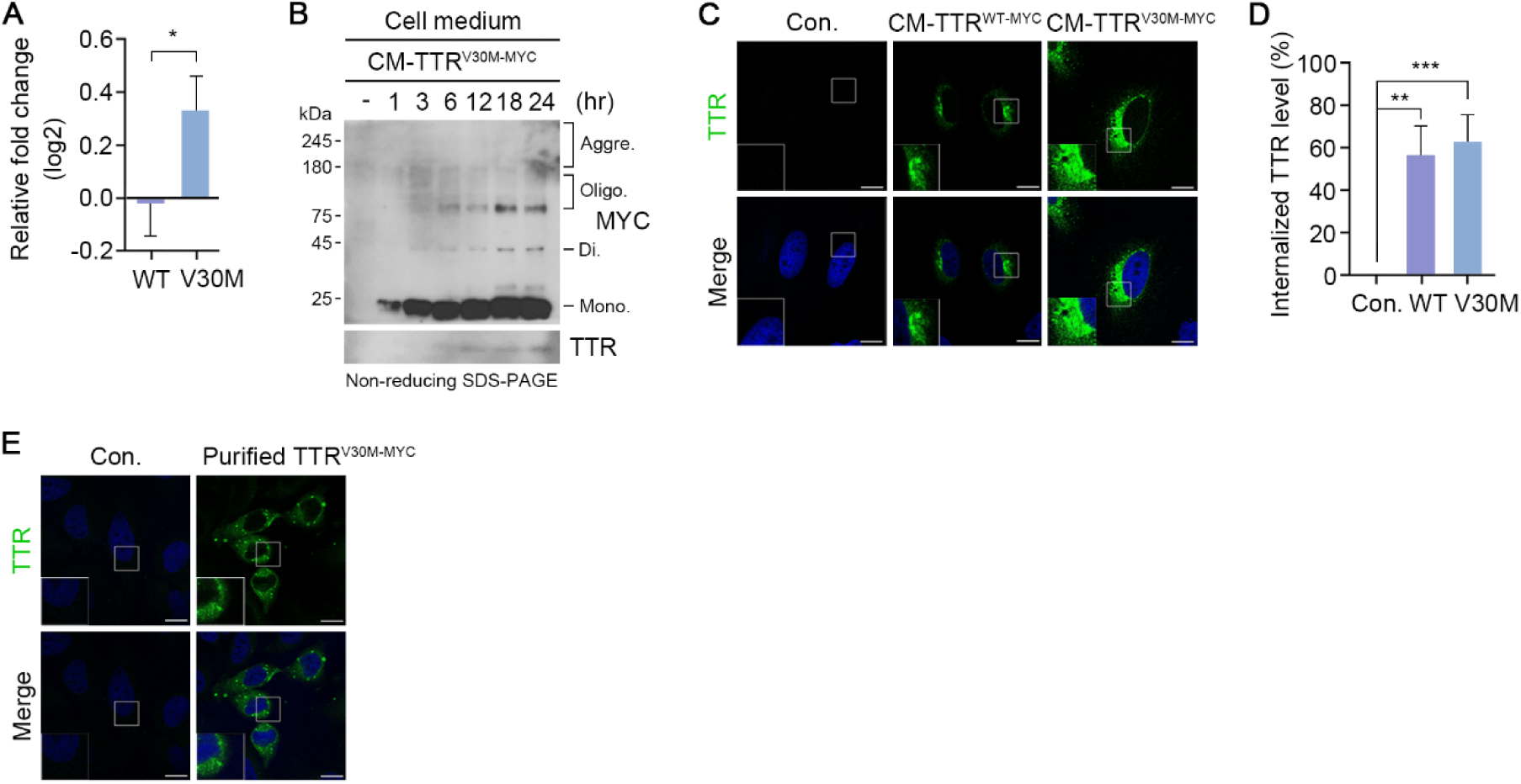
Amyloidogenic TTR is internalized into the cytosol. (**A**) Relative fold change of Fig. 1B. (**B**) *In vivo* oligomerization assay of TCA-precipitated media from HeLa cells treated with CM containing secreted TTR^V30M-MYC^ at the indicated time points. (**C**) Immunostaining analysis of HeLa cells treated with CM containing secreted WT TTR^MYC^ or TTR^V30M-MYC^ for 24 h. (**D**) Quantification of (C) (n = 3, biologically independent experiments each counting 50 cells). (**E**) Immunostaining analysis of HeLa cells incubated with purified Myc-tagged TTR^V30M^ proteins from HEK293T cells (0.5 μg/ml, 24 h). Scale bar represents 10μm. Values represent mean ± SD. Each n represents an independent biological replicate. Unpaired two-tailed Student’s *t*-test (A); one-way ANOVA (D); p* < 0.05, **p < 0.01, ***p < 0.001, ****p < 0.0001 and ns, not significant.

**Figure S2.**
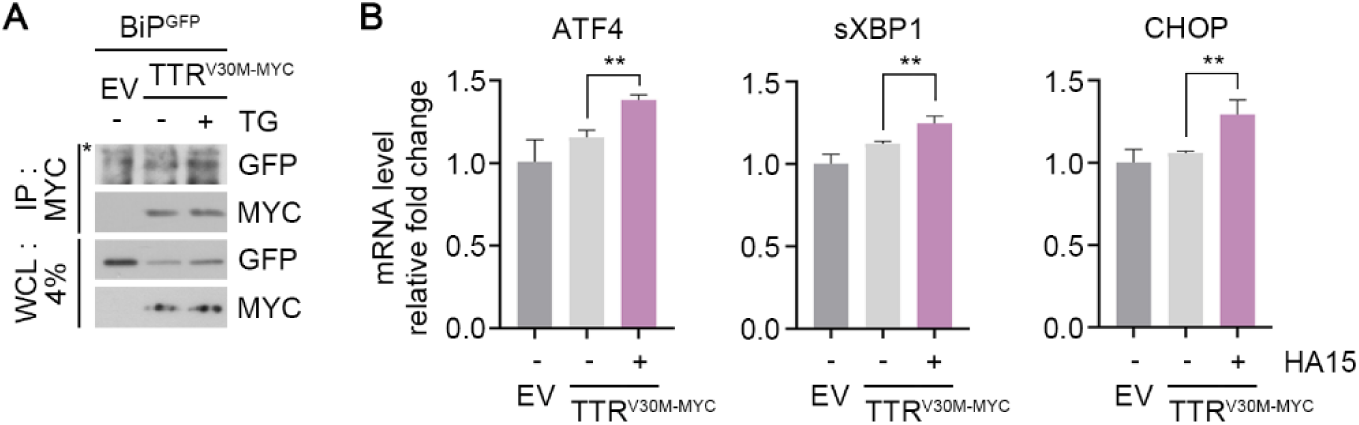
The ER-resident molecular chaperone BiP senses and regulates amyloidogenic TTR^V30M^. (**A**) CoIP assay in HEK293T cells co-transfected with TTR^V30M-MYC^ and BiP^GFP^, subsequently treated with thapsigarin (200 nM, 18 h). (**B**) The mRNA expression relative to GAPDH was measured by RT-qPCR. RNAs were isolated from HeLa cells transfected with TTR^V30M-MYC^, subsequently treated with HA15 (5 μM, 24 h) (n=3). Scale bar represents 10μm. Values represent mean ± SD. Each n represents an independent biological replicate. One-way ANOVA; p* < 0.05, **p < 0.01, ***p < 0.001, ****p < 0.0001 and ns, not significant.

**Figure S3.**
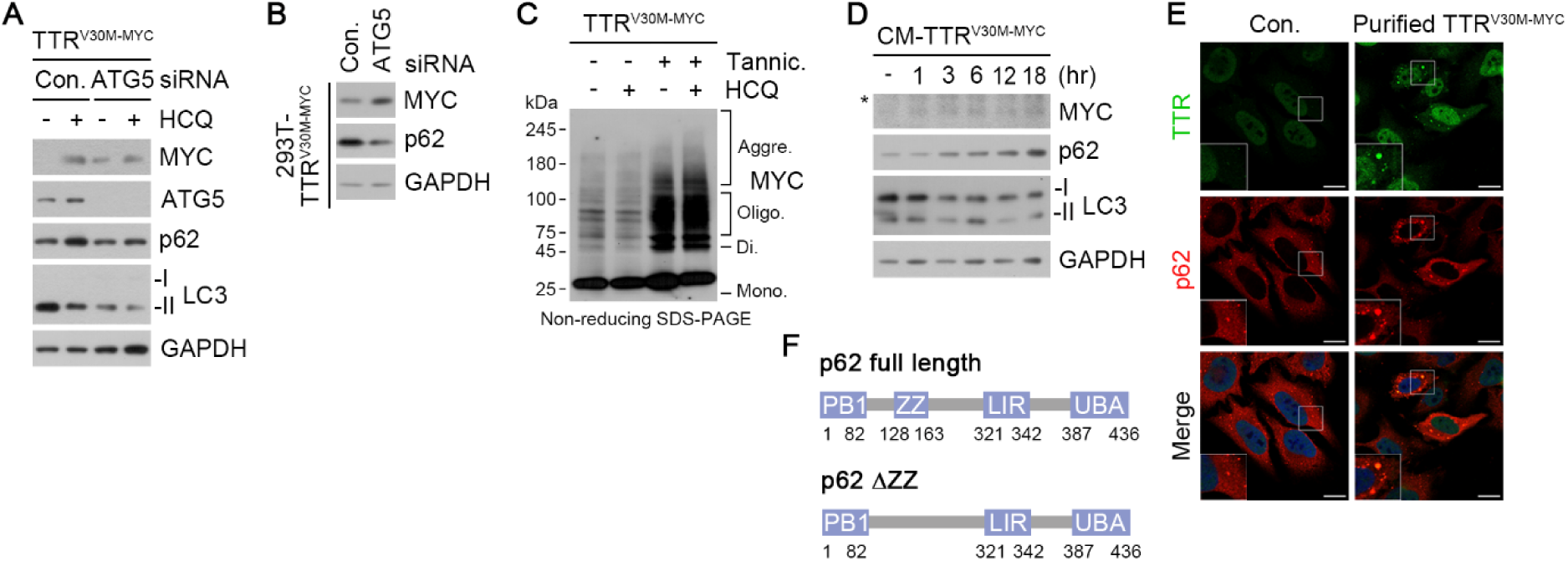
R-BiP bridges p62 and pathological TTR to promote lysosomal degradation. (**A**) Immunoblotting analysis of HEK293T cells co-transfected with TTR^V30M-MYC^ and siRNAs, subsequently treated with HCQ (10 μM, 24 h). (**B**) Immunoblotting analysis of TTR^V30M-MYC^ expressing 293T stable cell line transfected with siRNAs. (**C**) *In vivo* oligomerization assay of HeLa cells transfected with TTR^V30M-MYC^, subsequently treated with tannic acid (25 μM, 24 h) in the presence or absence of HCQ (10 μM, 24 h). (**D**) Immunoblotting analysis of HeLa cells treated with CM containing secreted TTR^V30M-MYC^ at the indicated time points. (**E**) Immunostaining analysis of HeLa cells incubated with purified Myc-tagged TTR^V30M^ proteins from HEK293T cells (0.5 μg/ml, 24 h). (**F**) Schematic illustration of full length p62 in comparison with ΔZZ p62. Scale bar represents 10μm. Asterisks (*) denote non-specific bands.

**Figure S4.**
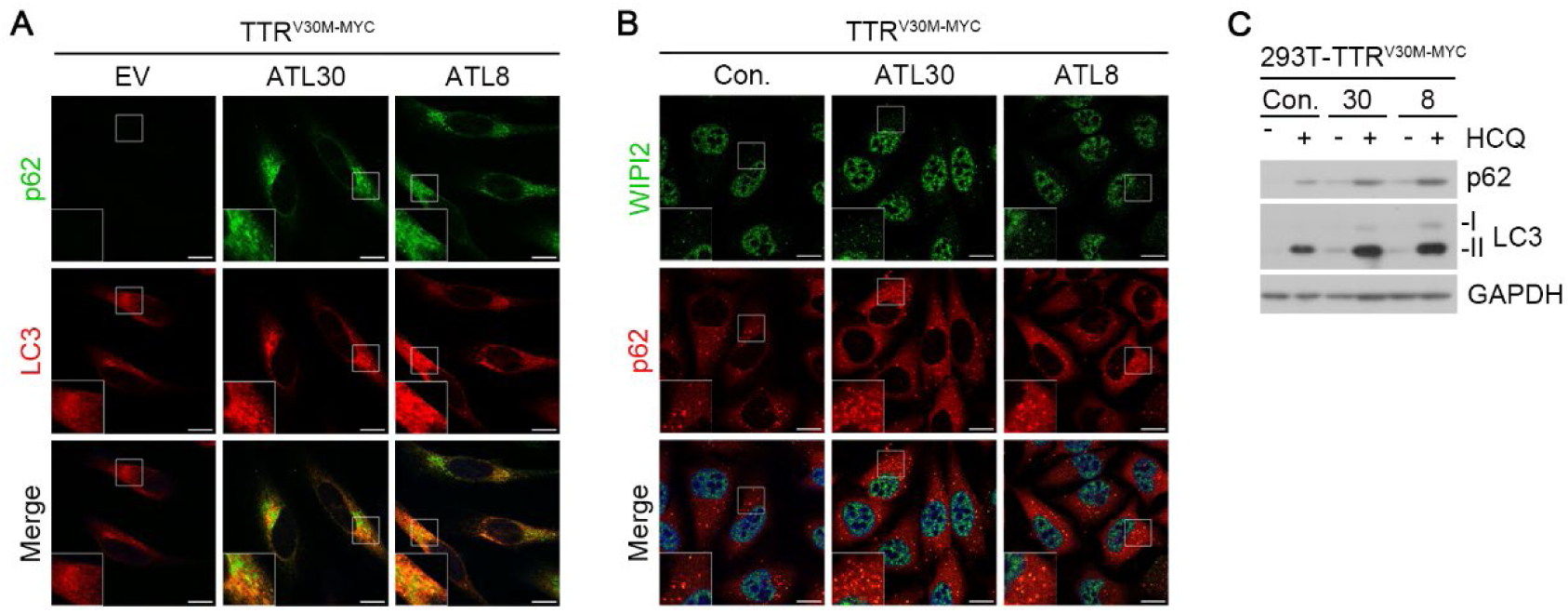
ATLs upregulates autophagic formation and flux in cells transiently or stably expressing TTR^V30M-MYC^. (**A**) Immunostaining analysis of HeLa cells transfected with TTR^V30M-MYC^, subsequently treated with ATL30 or ATL8 (1 μM, 24 h). (**B**) Identical to (A). (**C**) Immunoblotting analysis of TTR^V30M-MYC^ expressing 293T stable cell line treated with ATL30 or ATL8 (2.5 μM, 24 h) in the presence or absence of HCQ (10 µM, 24 h). Scale bar represents 10 μm.

**Figure S5.**
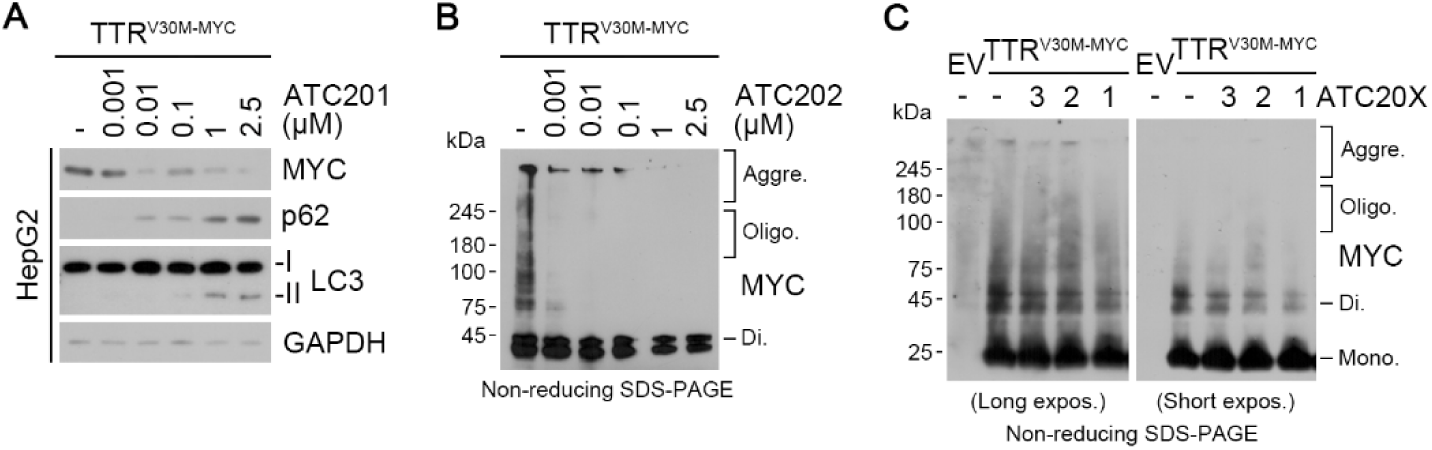
Autotacs degrade TTR. (**A**) Immunoblotting analysis of HepG2 cells transfected with TTR^V30M-MYC^, subsequently treated with ATC201 at the indicated concentrations (24 h). (**B**) *In vivo* oligomerization assay of HeLa cells transfected with TTR^V30M-MYC^, subsequently treated with ATC202 at the indicated concentrations (24 h). (**C**) *In vivo* oligomerization assay of HeLa cells transfected with TTR^V30M-MYC^, subsequently treated with the indicated compounds (1 μM, 24 h).

**Figure S6.**
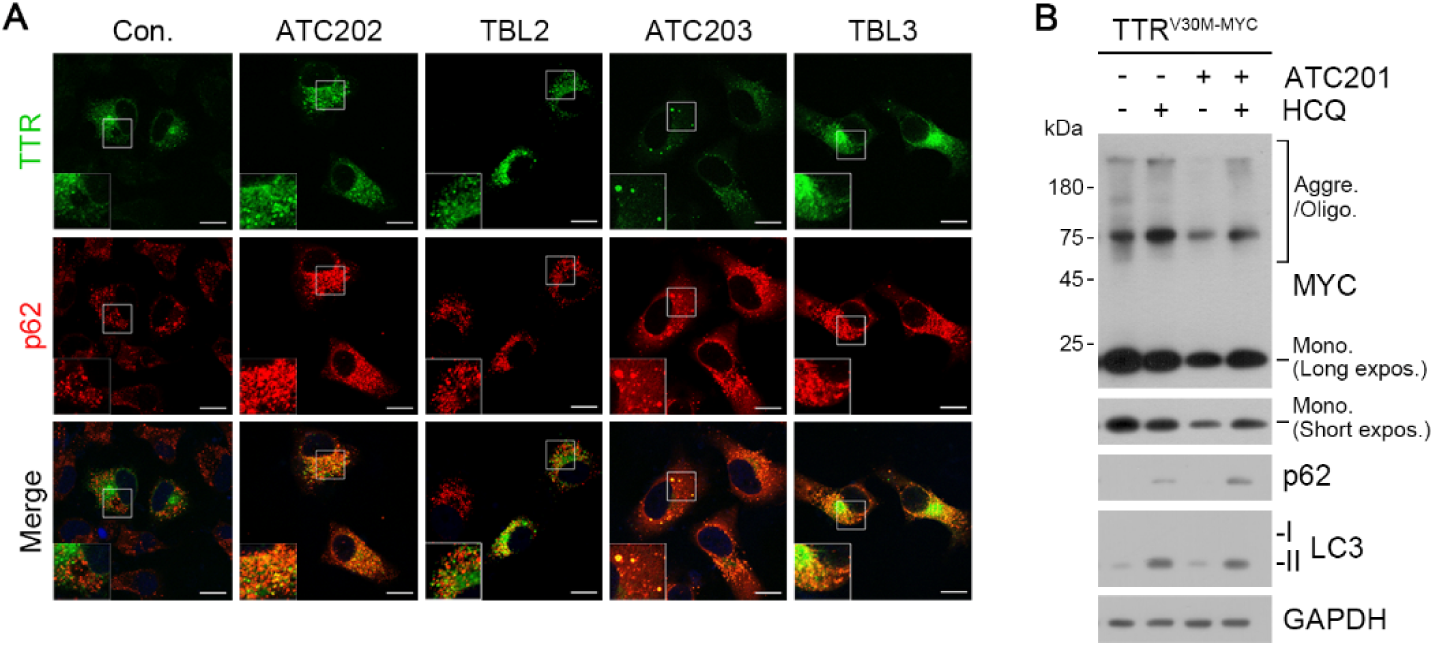
p62-depedent macroautophagic degradation of TTR via Autotacs. (**A**) Immunostaining analysis of HeLa cells transfected with TTR^V30M-MYC^, subsequently treated with the indicated compounds (1 μM, 24 h). (**B**) *In vivo* oligomerization assay of HeLa cells transfected with TTR^V30M-MYC^, subsequently treated with ATC201 (1 μM, 24 h) in the presence or absence of HCQ (10 μM, 24h). Scale bar represents 10 μm.

**Figure S7.**
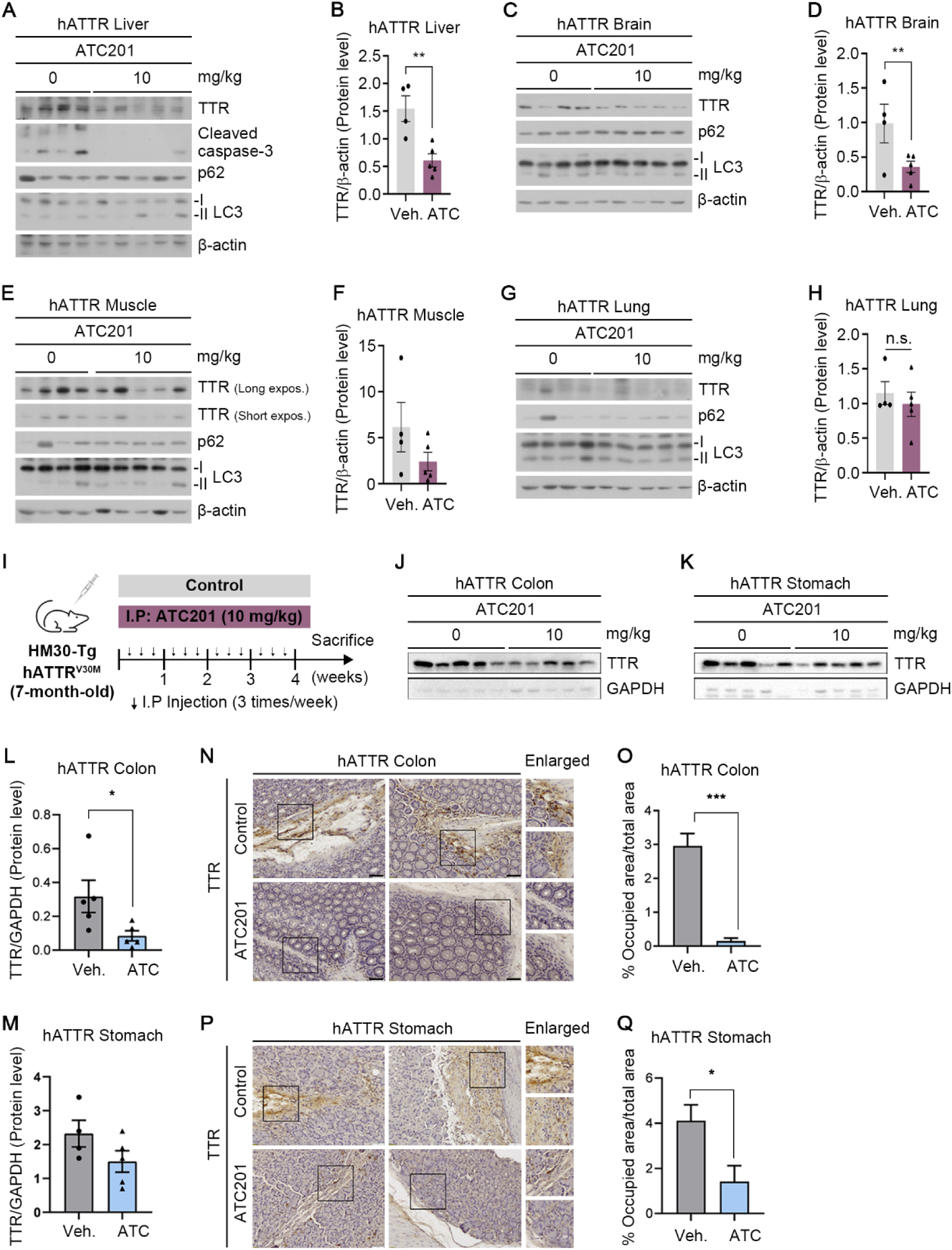
ATC201 mitigates TTR deposition in various tissues. (**A, C, E, G**) Immunoblotting analysis of liver, brain, muscle and lung tissues from hATTR mice injected with ATC201, as described in Fig. 7A. (**B, D, F, H**) Densitometry of TTR levels in (A, C, E, G), respectively (n = 9). (**I**) Schematic of hATTR^V30M^ murine model with injection timeline and details of ATC201. (**J-K**) Immunoblotting analysis of colon and stomach tissues from hATTR^V30M^ mice injected with ATC201, as described in Fig. 7A. (**L-M**) Quantification of TTR levels in (J) and (K), respectively (n = 10). (**N, P**) Immunohistochemistry analysis of colon and stomach tissues from hATTR^V30M^ mice injected with ATC201, as described in Fig. 7A. (**O, Q**) Quantification of (N) and (P), respectively. Scale bar represents 10 μm. Values represent mean ± S.E.M. Each n represents an independent biological replicate. Unpaired two-tailed Student’s *t*-test; p* < 0.05, **p < 0.01, ***p < 0.001, ****p < 0.0001 and ns, not significant.

**Figure S8.**
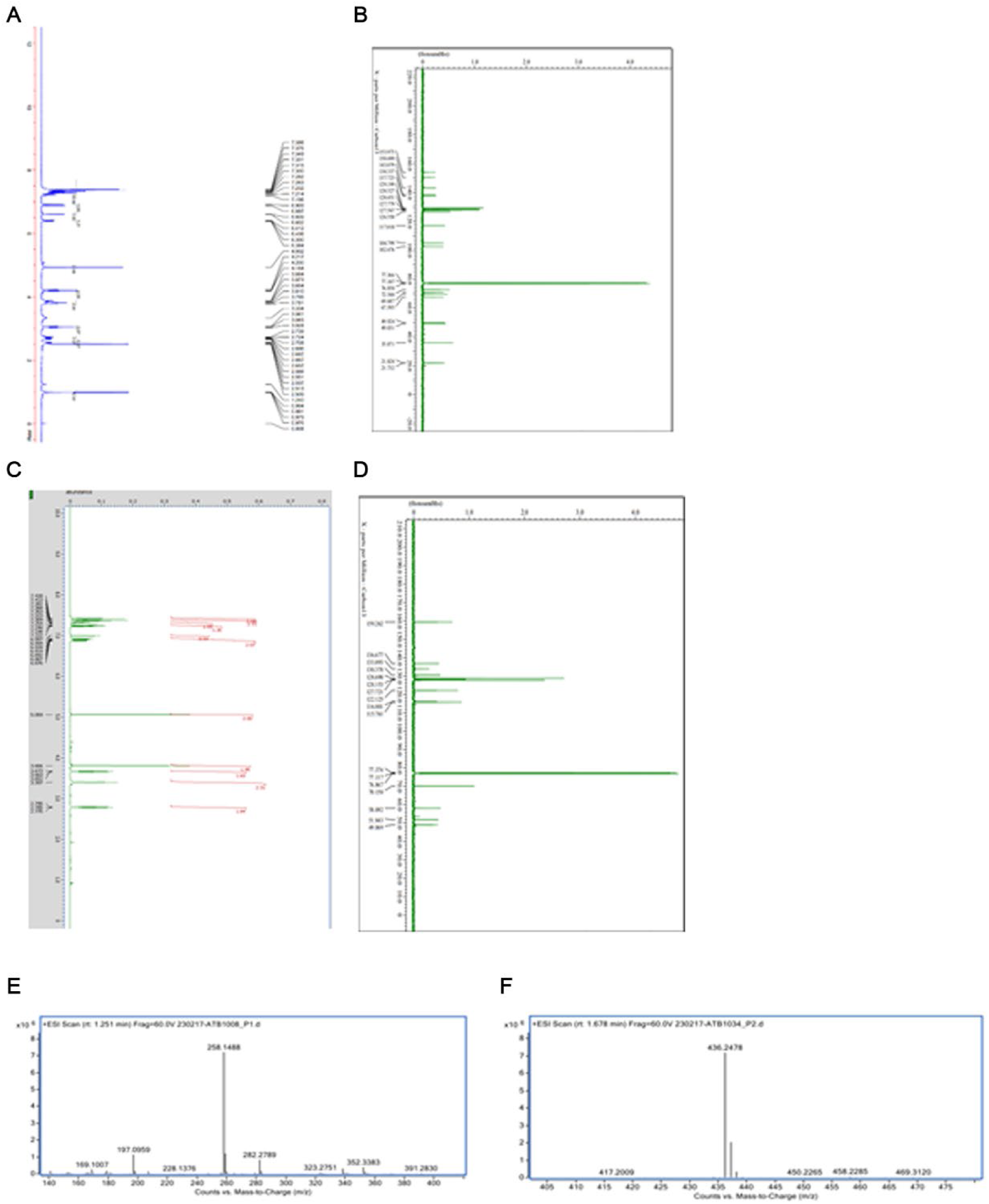
Spectroscopic and analytical charaterizaiton of ATLs and Autotacs. (**A**) 400 MHz ^1^H-NMR spectrum of (R)-1-(4-(benzyloxy)-3-phenethoxyphenoxy)-3-(isopropylamino) propan-2-ol (ATL30) in DMSO_d_6._ (**B**) 125 MHz ^13^C-NMR spectrum of (R)-1-(4-(benzyloxy)-3-phenethoxyphenoxy)-3-(isopropylamino) propan-2-ol (ATL30) in CDCl_3_. (**C**) 400 MHz ^1^H-NMR spectrum of 2-((3-(benzyloxy) benzyl) amino) ethan-1-ol (ATL08) in CDCl_3._ (**D**) 125 MHz ^13^C-NMR spectrum of 2-((3-(benzyloxy) benzyl) amino) ethan-1-ol (ATL08) in CDCl_3_. (**E**) HRMS data of (R)-1-(4-(benzyloxy)-3-phenethoxyphenoxy)-3-(isopropylamino) propan-2-ol (ATL30); HRMS Calcd m/z for C_27_H_33_NO_4_ [M+H]^+^ 436.2483 Found 436.2478. (**F**) HRMS data of 2-((3-(benzyloxy) benzyl) amino) ethan-1-ol (ATL08); HRMS Calcd m/z for C_16_H_19_NO_2_ [M+H]^+^ 258.1489 Found 258.1488.

## Supplementary tables

**Table S1.** MM-GBSA–based binding free energies (ΔG*_bind_*) of compounds against TTR and TTR–p62 complex.

| Compound ID | Receptor (PDB ID) | $\Delta G_{bind}$ (kcal/mol) |
| --- | --- | --- |
| Curcumin | TTR (PDB: 4PME) | -45.36 |
| ATC201 | TTR (PDB: 4PME) | -62.11 |
| ATC201 | TTR-p62 complex | -71.26 |

The $\Delta G_{bind}$ values were calculated using the MM-GBSA method to predict the affinity of the compounds (Curcumin and ATC201) for their respective receptors.

**Table S2.** PK parameters of ATC201 (single injection) in ICR mice.

| Route | IP | PO | IV |
| --- | --- | --- | --- |
| Dose/Unit (mpk) | 10 | 10 | 1 |
| $T_{1/2}$ (h) | 2.91 | | 0.73 |
| $T_{max}$ (h) | 0.25 | 0.67 | 0.083 |
| $C_{max}$ (ng/ml) | 2092.4 | 7.7 | 5602 |
| $AUC_{last}$ (h*ng/ml) | 2881.7 | 6.76 | 1607.2 |
| F (%) |  | 0.042 |  |

Mice received ATC201 via intravenous (IV), intraperitoneal (IP), or oral (PO) routes at the indicated doses (mg/kg). PK parameters include terminal half-life ( $T_{1/2}$ ), time to maximum concentration ( $T_{max}$ ), maximum plasma concentration ( $C_{max}$ ), area under the plasma concentration-time curve to the last measurable point ( $AUC_{last}$ ), and oral bioavailability (F, %).

## Supplementary methods

### Chemical synthesis and analytical data of Nt-Arg-mimicking compounds.

1H-NMR and 13C-NMR spectra were recorded on Bruker Advance III 500 MHz and 400 MHz and TMS was used as an internal standard. LCMS was taken on a quadrupole Mass Spectrometer on Agilent 1260 HPLC and 6120MSD (Column: C18 (50 × 4.6 mm, 5 μm) operating in ES (+) or (-) ionization mode; T = 30 °C; flow rate = 1.5 mL/min; detected wavelength: 254 nm. LC-HRMS was taken by 3 methods. One was taken on Agilent 6550 iFunnel Q-TOF (Column: ZORBAX RRHD SB-C18 (80Å, 2.1x100mm 1.8μm)) operating in ES (+) or (-) ionization mode; T = 25 °C; flow rate = 0.5 mL/min; detected wavelength: 254 nm. Another was taken on Agilent G6520 Q-TOF (Column: Agilent EC-C18 (4.6x50mm, 4.0um)) operating in ES (+) or (-) ionization mode; T = 25 °C; flow rate = 1.5 mL/min; detected wavelength: 220nm and 254 nm. The other was taken on Agilent G6520 Q-TOF (Column: Xbridege C18 (4.6x50mm, 5.0um)) operating in ES (+) or (-) ionization mode; T = 25 °C; flow rate = 1.5 mL/min; detected wavelength: 220nm and 254 nm.

### Scheme 1. 2-((3-(benzyloxy) benzyl) amino) ethan-1-ol (ATL08)

#### 1.1 Synthesis of 3-(benzyloxy) benzaldehyde (1-1)

To a solution of 1 (3-hydroxybenzaldehyde) (0.5 g, 4.09 mmol) and benzyl bromide (1.05 g, 6.14 mmol) in DMF (9 mL) was add K2CO3 (0.85 g, 6.14 mmol) at 25 °C. The mixture was stirred for overnight at 60 °C. The reaction was poured into water, then was filtered to give 1-1 (3-(benzyloxy) benzaldehyde, 0.72 g, 83 % yield). 1H-NMR (DMSO_d6, 400 MHz) δ (ppm) 7.55-7.51 (m, 3H), 7.48-7.46 (m, 2H), 7.40 (t, 2H, J = 5.6 Hz), 7.37-7.33 (m, 2H), 5.20 (s, 2H)

#### 1.2 Synthesis of 2-((3-(benzyloxy) benzyl) amino) ethan-1-ol (ATL08)

To a solution of 1-1 (3-(benzyloxy) benzaldehyde, 0.7 g, 3.30 mmol) were added 2-aminoethanol (0.18 g, 2.97 mmol) and AcOH (0.5 ml) in MeOH (13 mL). The mixture was stirred for overnight at 50 °C. The mixture was cooled to r.t and NaBH3CN (0.41 g, 6.60 mmol) was added slowly at R.T. Then the mixture was stirred for overnight at R.T. The reaction was evaporated in vacuo. Water was poured into the mixture and extracted with DCM. The organic layer was dried over MgSO4, filtered and concentrated. The crude was purified by column chromatography (DCM/MeOH=20/1∼10/1) to give ATL08 (2-((3-(benzyloxy) benzyl) amino) ethan-1-ol, 0.3 g, 30 % yield) as pale-yellow oil. 1H-NMR (CDCl3, 500 MHz): δ (ppm) 9.42 (br s, 2H), 7.39 (d, 2H, J = 7.0 Hz), 7.33 (t, 1H, J = 7.0 Hz), 7.30-7.24 (m, 3H), 7.10 (d, 1H, J = 6.0 Hz), 6.95 (d, 1H, J = 8.5 Hz), 5.06 (s, 2H), 4.13 (br s, 2H), 3.89 (br s, 2H), 2.95 (br s, 2H); 13C-NMR (CDCl3, 125 MHz): δ (ppm) 159.26, 136.68, 133.89, 130.37, 128.69, 128.15, 127.72, 122.13, 116.00, 115.76, 70.16, 58.09, 51.84, 49.07; HRMS Calcd m/z for C16H19NO2 [M+H]+ 258.1489 Found 258.1488

### Scheme 2. (R)-1-(4-(benzyloxy)-3-phenethoxyphenoxy)-3-(isopropylamino) propan-2-ol (ATL30)

#### 2.1 Synthesis of 4-(benzyloxy)-3-hydroxybenzaldehyde (2-1)

To a solution of 2 (3,4-dihydroxybenzaldehyde, 20.0 g, 145 mmol, 1.0 eq) and (bromomethyl)benzene (24.8 g, 145 mmol, 1.0 eq) in ACN (400 mL) was add NaHCO3 (14.6 g, 174 mmol, 1.2 eq) at 25 °C. The mixture was stirred overnight at 80 °C. The reaction was concentrated. The residue was quenched with 1N HCl and extracted with EA. The organic layer was washed with brine, dried over Na2SO4, filtered and concentrated. The residue was purified by silica gel, eluted with EA/PE (20:1∼10:1) to afford 2-1 (4-(benzyloxy)-3-hydroxybenzaldehyde, 10.0 g, yield: 30.3%). (TLC: PE/EA=5/1, Rf=0.6) 1H-NMR (DMSO_d6, 400 MHz): δ (ppm) 9.76 (s, 1H), 9.66 (s, 1H), 7.49-7.48 (m, 2H), 7.42-7.34 (m, 4H), 7.29 (d, J = 2.0 Hz, 1H), 7.20 (d, J = 8.4 Hz, 1H), 5.23 (s, 2H).

#### 2.2 Synthesis of 4-(benzyloxy)-3-phenethoxybenzaldehyde (2-2)

To a solution of 2 (4-(benzyloxy)-3-hydroxybenzaldehyde, 10.0 g, 43.9 mmol, 1.0 eq) and (2-bromoethyl) benzene (9.71 g, 52.6 mmol, 1.2 eq) in DMF (100 mL) was added Cs2CO3 (43.0 g, 132 mmol, 3.0 eq). The mixture was stirred at 80 °C overnight. The mixture was added water and extracted with EA. The organic layer was washed with brine, dried over Na2SO4, filtered and concentrated. The residue was purified by silica gel, eluted with EA/PE (20:1∼10:1) to afford 2-2 (4-(benzyloxy)-3-phenethoxybenzaldehyde, 3.7 g, yield: 25.3%). (TLC: PE/EA=3/1, Rf=0.6) 1H-NMR (CDCl3, 400 MHz): δ (ppm) 9.83 (s, 1H), 7.46-7.25 (m, 12H), 7.02 (d, J = 7.6 Hz, 1H), 5.22 (s, 2H), 4.31 (t, J = 6.8 Hz, 2H), 3.18 (t, J = 6.8 Hz, 2H).

#### 2.3 Synthesis of 4-(benzyloxy)-3-phenethoxyphenol (2-3)

To a solution of 2-2 (4-(benzyloxy)-3-phenethoxybenzaldehyde, 3.70 g, 11.1 mmol, 1.0 eq) in DCM (40 mL) was added mCPBA (2.90 g, 16.7 mmol, 1.5 eq) in portions. The mixture was stirred at rt for 2 hrs. The mixture was washed with saturated NaHCO3 solution, and concentrated. The mixture was dissolved in MeOH (25 mL) and added 5N KOH (2.5 mL, 12.3 mmol, 1.1 eq). The mixture was stirred at rt for 1 hour. The mixture was added ice water and filtered. The solid was concentrated to afford 2-3 (4-(benzyloxy)-3-phenethoxyphenol, 3.4 g, yield: 95.5%). (TLC: PE/EA=3/1, Rf=0.3) 1H-NMR (DMSO_d6, 400 MHz): δ (ppm) 9.01 (s, 1H), 7.37-7.21 (m, 10H), 6.80 (d, J = 8.8 Hz, 1H), 6.43 (s, 1H), 6.22 (dd, J = 2.4, 8.4 Hz, 1H), 4.87 (s, 2H), 4.14 (t, J = 6.8 Hz, 2H), 3.03 (t, J = 6.4 Hz, 2H).

#### 2.4 Synthesis of (R)-2-((4-(benzyloxy)-3-phenethoxyphenoxy) methyl) oxirane (2-4)

To a solution of 2-3 (4-(benzyloxy)-3-phenethoxyphenol, 3.4 g, 10.6 mmol, 1.0 eq) in EtOH (50 mL) was added KOH (0.7 g, 12.8 mmol, 1.2 eq) and H2O (5 mL). The mixture was added (R)-2-(chloromethyl) oxirane (2.9 g, 31.9 mmol, 3.0 eq). The mixture was stirred at 30 °C overnight. The mixture was added water and filtered. The solid was concentrated to give 2-4 ((R)-2-((4-(benzyloxy)-3-phenethoxyphenoxy) methyl) oxirane, 3.6 g, yield: 90%). (TLC: PE/EA=3/1, Rf=0.6) 1H-NMR (DMSO_d6, 400 MHz): δ (ppm) 7.41-7.19 (m, 10H), 6.90 (d, J = 8.8 Hz, 1H), 6.65 (d, J = 2.8 Hz, 1H), 6.42 (dd, J = 2.4, 8.8 Hz, 1H), 4.93 (s, 2H), 4.26-4.18 (m, 3H), 3.77-3.72 (m, 1H), 3.30-3.27 (m, 1H), 3.04 (t, J = 6.4 Hz, 2H), 2.82 (t, J = 4.8 Hz, 1H), 2.69-2.67 (m, 1H).

#### 2.5 Synthesis of (R)-1-(4-(benzyloxy)-3-phenethoxyphenoxy)-3-(isopropylamino) propan-2-ol (ATL1)

The mixture of 2-4 ((R)-2-((4-(benzyloxy)-3-phenethoxyphenoxy) methyl) oxirane, 3.6 g, 9.6 mmol, 1.0 eq) and propan-2-amine (2.8 g, 47.9 mmol, 5.0 eq) in MeOH (100 mL) was stirred overnight at 50 °C. The reaction mixture was concentrated and purified by chromatography (DCM/MeOH=15/1) to give ATL08 ((R)-1-(4-(benzyloxy)-3-phenethoxyphenoxy)-3-(isopropylamino) propan-2-ol, 1.0 g, yield: 24.0%). (TLC: DCM/MeOH=15/1, Rf=0.2)

1H-NMR (DMSO_d6, 400 MHz): δ (ppm) 7.38-7.19 (m, 10H), 6.90 (d, J = 8.8 Hz, 1H), 6.60 (d, J = 2.8 Hz, 1H), 6.40 (dd, J = 2.4, 8.8 Hz, 1H), 4.93 (s, 2H), 4.20 (t, J = 6.8 Hz, 2H), 3.88-3.78 (m, 3H), 3.04 (t, J = 6.4 Hz, 2H), 2.73-2.65 (m, 2H), 2.56-2.50 (m, 1H), 0.99 (dd, J = 1.2, 6.0 Hz, 6H); 13C-NMR (CDCl3, 125 MHz): δ (ppm) 153.97, 150.49, 143.08, 138.34, 137.72, 129.20, 128.53, 128.45, 127.78, 127.57, 126.55, 117.02, 104.80, 102.48, 72.55, 69.69, 67.60, 49.92, 49.03, 35.87, 21.83, 21.73; HRMS Calcd m/z for C27H33NO4 [M+H]+ 436.2483 Found 436.2478

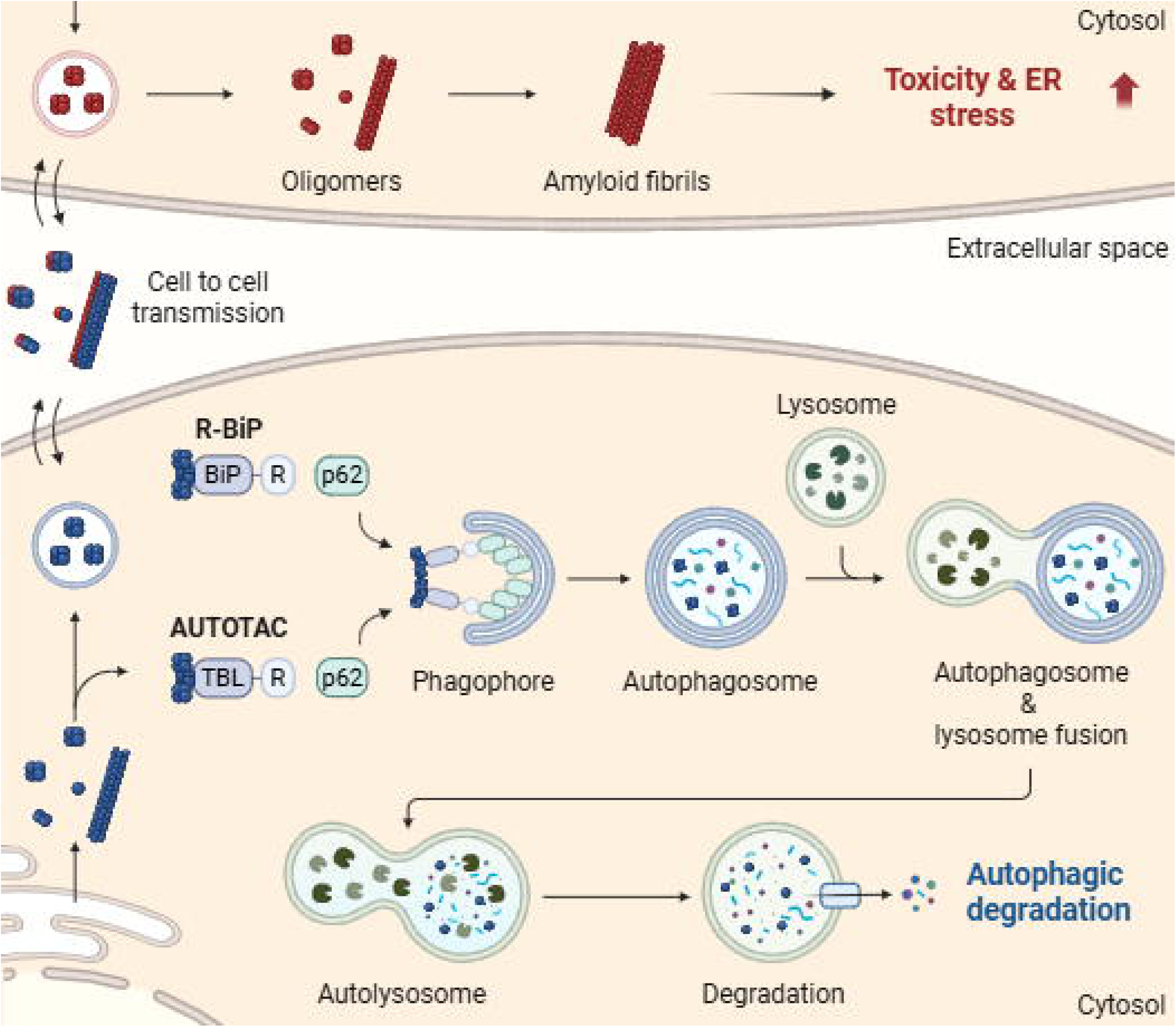

